# Chinmo is a novel regulator of differential response to Hippo signaling within a single developing organ

**DOI:** 10.1101/2025.05.02.651944

**Authors:** Shreeharsha Tarikere, Tarun Kumar, Guillem Ylla, Ana A. Stratan, Cassandra G. Extavour

## Abstract

Orchestrated control of proliferation of multiple cell types is essential for building a healthy organ. Here we use larval ovary development in *Drosophila melanogaster* as a model to understand the homeostasis of somatic and germ line cells in the formation of the female adult reproductive organ. We previously showed that the highly conserved Hippo signaling pathway regulates proliferation of both germ line and somatic cells in the *D. melanogaster* larval ovary. Response to Hippo signaling appeared to be mediated by different genetic mechanisms in germ line and soma, but the mechanism allowing distinct responses to the same signaling pathway remained unknown. Here we perform cell type-specific RNA-Seq of isolated germ line and somatic cells from the developing ovary at multiple time points spanning the formation of germ line stem cell niches, in *hpo* and *yki* knockdown conditions. Applying network analysis to these data revealed a novel regulator of ovarian development, the transcription factor *chinmo*. Subsequent experimental validation suggested that *chinmo* acts as a key germ line-specific translator of Hippo signaling in the developing ovary, potentially allowing the Hippo signal to be transduced in cell type-specific ways in germ line and somatic cells within the same organ.

## INTRODUCTION

Animal organs are not homogeneous masses of cells. Instead, they are composed of multiple cell types that contribute to the same organ, but have distinct gene expression profiles, morphologies, behaviours, and functions. In most organ systems, all of these distinct cell types are somatic cells derived from the same germ layer (Barresi and Gilbert, 2023). An important exception to this rule is found in the gonads (ovaries and testes), which contain both somatic cells of mesodermal origin, and cells of the germ line, destined to make eggs and sperm (Extavour, 2007). Gonads are typically assembled after the specification of germ layers and of the germ line, but before gametogenesis begins. During gonadogenesis, somatic and germ cells proliferate, exchange neighbors, change their shapes and gene expression profiles, and create the final functional and morphological organization of the gonad. As for other systems of organogenesis, cell-cell signaling is crucial for this process. However, each unique cell type within the developing organ often has a distinct, cell type-appropriate response to the same incoming signals, allowing correct organ formation and function. Here we use the *Drosophila melanogaster* (fruit fly) ovary as a study system to elucidate how different cell types within the same organ can differently transduce the same signaling inputs during development.

A single adult ovary in *D. melanogaster* is made up of an estimated 26 distinct cell types (Rust et al., 2020) that are assembled into 16-23 tubular egg producing subunits called ovarioles (Büning, 1994), making it an excellent model to study the regulation of organ size and subunit number (Kumar et al., 2020; Sarikaya and Extavour, 2015). Ovariole number varies by up to four orders of magnitude across insect species (Church et al., 2021). and in some insects, including Drosophilids, it is positively correlated with egg-laying and therefore fitness (Church et al., 2021; David, 1970). In *Drosophila* species, ovariole number is established in the larval stages and determined by the number of stacks (called terminal filaments: TFs) of a specific type of somatic cell (called terminal filament cells: TFCs) (Green and Extavour, 2012; Hodin, 2009; Honěk, 1993; King et al., 1968; Markow and O’Grady, 2007; Sarikaya et al., 2019). The number of these stacks of cells, in turn, is determined by the number and stacking dynamics of the component TFCs. We previously showed that TFC proliferation, and therefore ovariole number, is regulated by the Hippo (Hpo) signaling pathway, which affects both TFC and germ cell numbers (Sarikaya and Extavour, 2015).

First discovered as a network of tumor overgrowth factors in *D. melanogaster* (Dong et al., 2007; Tapon et al., 2002; Wu et al., 2003; Xu et al., 1995), the Hpo signaling pathway regulates organ size, growth, cell proliferation and cell fate determination in animals. In the developing fruit fly ovary, this pathway regulates proliferation both cell-autonomously (based on cell-intrinsic Hpo signaling activity) and non-cell autonomously (activity in one cell can affect proliferation in a different cell) (Sarikaya and Extavour, 2015). Regulation of cell proliferation in both the soma and the germ line is essential for the normal development of the ovary, including in determining species-specific ovariole number (Green and Extavour, 2012; Sarikaya et al., 2012). Members of the Hippo signaling pathway are expressed ubiquitously across multiple distinct ovarian cell types that nevertheless have different proliferation dynamics (Sarikaya and Extavour, 2015). This begs the question of how differential regulation of Hpo signaling may be achieved despite the potential for ubiquitous activation.

The canonical Hippo signaling pathway functions through regulating shuttling of the transcriptional activator Yorkie (Yki) in and out of the nucleus. This shuttling is regulated by the function of the serine-threonine kinase Hpo in conjunction with a cascade of kinases that ends with the phosphorylation of Yki, preventing its nuclear localization (Huang et al., 2005; Oh and Irvine, 2008). Under this model, unphosphorylated Yki enters the nucleus to form tissue-specific transcriptional regulatory complexes (Slattery et al., 2013) with loading proteins like Scalloped (Sd) and Homothorax (Hth). These transcription cofactor complexes bind to DNA and activate a wide range of target genes, including many necessary for growth and proliferation (Manning et al., 2024; Peng et al., 2009; Wu et al., 2008; Zhang et al., 2008; Zhao et al., 2008). Phosphorylation of Yki by other kinases, including Nemo-like kinase and AMP-activated protein kinase, can either prevent or enhance Yki’s targeting by the canonical Hpo kinases, allowing a Hpo-independent mode of Yki regulation (Moon et al., 2017; Wang et al., 2015). Furthermore, changes in the binding dynamics of Sd, a coactivator of Yki, adds another layer of Yki regulation (Manning et al., 2024). Lastly, mechanical stress and proximity of cell membranes during cell growth and proliferation can also regulate Hpo signaling activity (Zhao et al., 2007). Across animals, the Hpo pathway regulates multiple cellular processes *via* Yki target genes, including cell size, cell proliferation, adhesion and apoptosis (Zheng and Pan, 2019), while interacting with other signaling pathways such as JAK-STAT, Notch, Hedgehog and EGFR (Besse et al., 2005; Kumar et al., 2020; Matsuoka et al., 2013; Sarikaya and Extavour, 2015; Song et al., 2007; Yatsenko and Shcherbata, 2021).

Previously, we showed that Hpo pathway activity regulates cell proliferation in multiple *D. melanogaster* ovarian cell types, which in turn can determine the numbers of TFCs, TFs and ultimately ovarioles (Sarikaya and Extavour, 2015). RNAi-mediated knockdown of *hpo* or *warts* in somatic cells increased TF and TFC number, while RNAi against *yki* decreased the numbers of both cell types (Sarikaya and Extavour, 2015). In this context, we demonstrated that the Hpo pathway interacts with the EGFR pathway to regulate numbers of another somatic cell type, called intermingled cells (Sarikaya and Extavour, 2015) and with JAK-STAT signaling to regulate TFC proliferation and the homeostasis between intermingled cells and germ cells (Sarikaya and Extavour, 2015). Furthermore, our RNAi modifier screen in a *hpo* knockdown background in the larval ovary identified a gene interaction network implicating all known conserved animal signaling pathways in regulating *D. melanogaster* ovary development and function, including establishment of correct ovariole number (Kumar et al., 2020).

To our surprise, our previous study showed that in contrast to somatic cells, *hpo* knockdown in the germ line did not affect germ cell number (Sarikaya and Extavour, 2015). However, *yki* knockdown in the germ line reduced germ cell number significantly, as it did in somatic cells. This result suggested the existence of a Hpo-independent mechanism of Yki activity in the germ cells (Sarikaya and Extavour, 2015). However, a mechanistic explanation for the differential response of germ cells and somatic cells to these Hpo signaling components remains lacking. Given the variation in outcomes of Hpo pathway abrogation in the germ cells and the somatic cells, we sought to use cell type-specific transcriptional profiling to identify distinct gene regulatory responses to *hpo* or *yki* activity in soma versus germ line. Here we present the results of this approach, wherein we analyze stage- and tissue-specific transcriptomic datasets generated in three distinct genetic backgrounds to assess the role of Hpo signaling and its interacting pathways in the cell types of the developing *D. melanogaster* larval ovary. We reveal the transcription factor Chinmo (Zhu et al., 2006) as a putative novel regulator of germ cell-specific Hpo-dependent activity and provide experimental validation of this regulatory interaction in the developing ovary. Further, we provide evidence that *chinmo* regulates the previously observed (Sarikaya and Extavour, 2015) non-autonomous effect that Hpo activity in germ cells exerts on neighboring somatic cells. Thus, our results show how variation in the expression of cell type-specific targets can produce distinct proliferation response outcomes to the same signaling inputs within a developing tissue.

## RESULTS

### *hpo* and *yki* knockdown in somatic and germ line tissues

*D. melanogaster* ovaries are made up of subunits called ovarioles (Figure 1A), whose number is established at the third larval instar stage, 72-120 hours after egg laying (hAEL) at 25°C (Figure 1B) (King, 1970). The number of ovarioles in the adult corresponds to the number of TFs formed in the larval ovary (Figure 1B), because TFs are neither formed nor destroyed during the pupal phase that separates larva from adult (King, 1970). The Hpo signaling pathway regulates TF number by determining the number of TFCs available to assemble into TFs (Sarikaya and Extavour, 2015). We reasoned that the genetic mechanisms of Hpo regulation of TF number might be revealed by determining the responses of germ cells and somatic cells to Hpo signaling activity during TF formation (Sarikaya and Extavour, 2015). We therefore selected larvae over three stages of early ovary morphogenesis as follows: (1) before TF formation (Early = 72 hAEL; Figure 1Bi); (2) during TF formation (Middle (mid) = 96 hAEL; Figure 1Bii); and (3) after TF formation (Late = 120 hAEL; Figure 1 Biii). We expressed GFP either in the somatic cells (red cells in Figure 1C) or in the germ cells (blue cells in Figure 1C) using cell type-specific GAL4 drivers (Brand and Perrimon, 1993) and isolated somatic and germ cells respectively using Fluorescence-Activated Cell Sorting (FACS) (Supplemental Figure S1).

**Figure 1:**
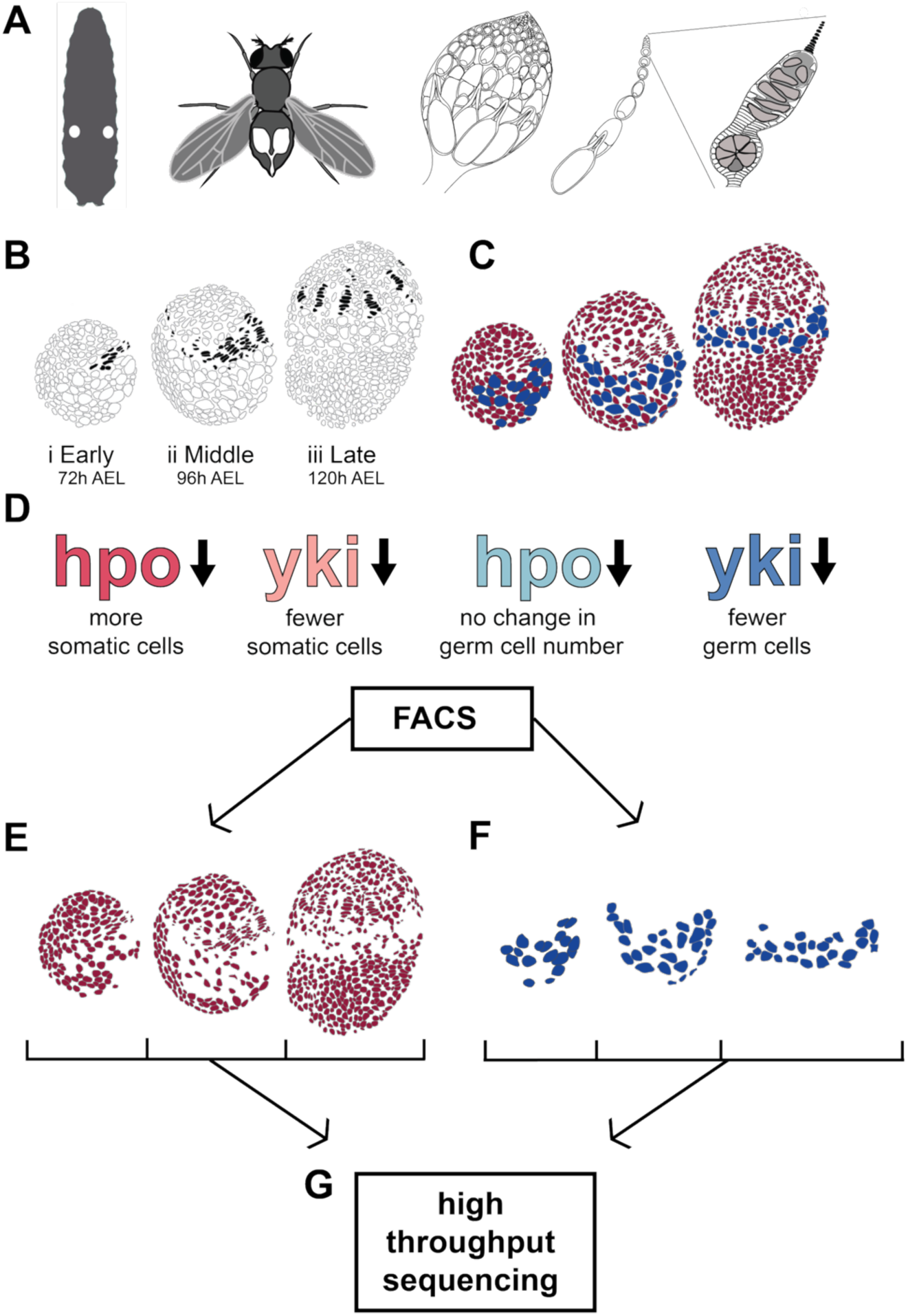
Experimental design to generate stage-specific transcriptomes of somatic and germ line larval ovarian cells under wild type, *hpo* and *yki* loss of function conditions. (**A**) Left to right: Location of larval ovaries (indicated as white circles in the larva); location of adult ovaries in female abdomen (indicated in white within the abdomen); a single adult ovary; a single ovariole; a single germarium with terminal filaments in black. (**B**) The three developmental stages during terminal filament (in black) formation studied herein. (**C**) Somatic (red) and germ line (blue) cells were marked for FACS sorting using *bab:GAL4* (soma) and *nanos:GAL4* (germ line) drivers as described in Materials and Methods. (**D**) Summary of the tissue-specific RNAi knockdown treatments assessed at each stage in somatic and germ line cells. Note that *hpo[RNAi]* induces no significant change in germ cell number under the *nanos:GAL4* driver relative to both parental controls (Sarikaya and Extavour, 2015 and Figure 5D). GFP labelled somatic cells (**E**) and germ cells (**F**) were sorted using FACS at the three developing stages of larval ovaries. Data from early-stage germ cells (grey), were not used for analysis; see Materials and Methods. (**G**) Sorted ovarian cell types were processed for mRNA extraction, cDNA library preparation and high throughput sequencing. h AEL = hours after egg laying.

We previously established the gene expression profiles of wild type somatic and germ cells at these larval stages, and reported dynamic changes in gene expression during ovariole formation (Tarikere et al., 2021). In the present study, we used the UAS/GAL4 system (Brand and Perrimon, 1993) to knock down either *hpo* or *yki* in either somatic or germ cells across the three developmental stages described above, and then performed tissue-specific RNA-Seq to identify the transcriptional regulatory effects of these two main components of the Hpo signaling pathway on larval ovary development and morphogenesis (Figure 1D-G). We used RNAi lines that were validated in multiple ways, as previously described (Sarikaya et al. 2015). Briefly, in these RNAi conditions, (1) both Hpo and Yki immunohistochemistry showed reduced protein levels in larval ovaries; (2) the effect of both RNAi lines on germ cell numbers were the same as that of *hpo* and *yki* null alleles; (3) a null allele of *expanded*, an upstream regulator of the Hpo pathway, has the same effect on TFC, germ cell, and interstitial cell number as *hpo* RNAi; (4). two different *yki* RNAi lines had the same effect on germ cell number. We used the *hpo* and *yki* RNAi lines thus validated, as well as the same *nanos:GAL4* (germ line-specific) and *bab:GAL4* (soma-specific) lines used in our previous work (Sarikaya and Extavour, 2015), to perform the experiments described herein.

### The Hpo pathway regulates gene expression differently in the ovarian germ line and soma

Our RNA-Seq study produced 48 independent datasets (Supplemental Table S1) of gene expression in germ cells and somatic cells in two different knockdown conditions and at three developmental stages. To identify and understand the variation in gene expression across samples, we performed a principal component analysis (PCA). Our results showed that 51.5% of the variation across all datasets was explained by cell type (Figure 2A). The first principal component clearly separated somatic cells from germ cells (PC1 in Figure 2A). Cell type-specific gene expression included both genes with known roles in the germ line (e.g. *vasa* (Lasko and Ashburner, 1988) and *bruno* (Kim-Ha et al., 1995)) or in the somatic gonad (e.g. *traffic jam* (Li et al., 2003) and *bab2* (Godt and Laski, 1995)), as well as novel genes without such roles previously described (Supplemental Table S2). The second greatest impact on transcriptional variation (PC2) was developmental stage (Figure 2A).

**Figure 2:**
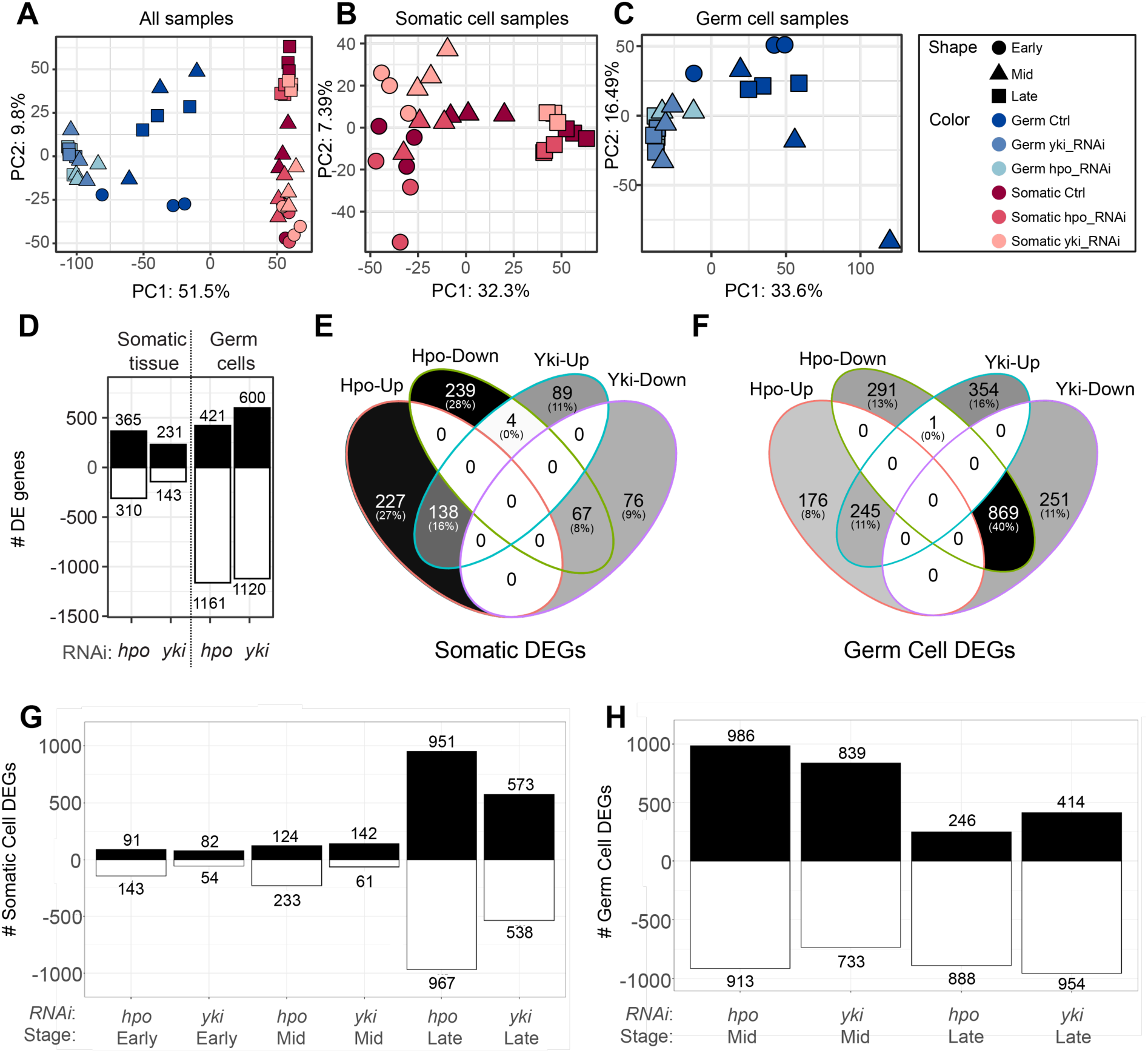
Distinct transcriptional impact of Hpo signaling on germ line and soma across ovarian development. **(A)** Principal Component Analysis (PCA) of all samples differentiates germ cells from somatic cells along PC1. (B) PCA of somatic transcriptomes showing clustering by time point in PC1, and the knockdown effect in PC2. **(C)** PCA of germ line transcriptomes in which control and knockdown libraries cluster separately, but the *hpo* and *yki* knockdowns cluster together. **(D)** Number of significantly upregulated and downregulated genes (adjusted p-value < 0.05) in each cell type with grouped stages when comparing each knockdown to the control library. For somatic cells **(E)** and germ cells **(F)** a Venn diagram shows the differentially up- and down-regulated genes under each knockdown, including number and percentage of genes in each intersection. **(G, H)** The number of significantly upregulated and downregulated genes (adjusted p-value < 0.01) per stage when comparing each knockdown to the control library, for somatic cells (G) and germ cells (H).

We then performed individual PCA analysis of somatic cells (Figure 2B) and germ cells (Figure 2C), which allowed us to better observe the effects of the two RNAi knockdown conditions on the sample clustering. In the somatic cells, most of the variation (32.3%) amongst the samples was attributable to developmental stage rather than to genotype (stages are indicated as distinct shapes along PC1 in Figure 2B), and most changes in gene expression occurred during the late developmental stage (Figure 2G). By contrast, in the germ cells, the samples clustered by genotype rather than by stage (colors along PC1 in Figure 2C). In other words, transcriptional changes under both conditions in the germ line were distributed across the developmental progression of the larval ovary (Figure 2H). The *hpo* and *yki* loss-of-function conditions clustered together independently of developmental stage, but far from the respective controls (Figure 2C). Along the PC2 axis, datasets were sorted by genotype, with both the *hpo* and *yki* loss-of-function treatments segregating in opposite directions relative to the control across all three developmental stages (colors along PC2 in Figure 2B).

These results suggest that *yki* and *hpo* regulate transcriptional activity in ovarian germ cells and somatic cells differently during larval development. In the soma, RNAi against each of the two genes had distinct and stage-specific impacts on the transcriptomic landscape. Notably, in somatic cells there were significantly more differentially expressed genes, both upregulated and downregulated, at the late larval stage compared to early and mid stages (Figure 2G). In contrast, loss of function of both genes had very similar effects on the transcriptomes of the germ line, and these regulatory impacts did not change significantly from mid to late stages of development (Figure 2H). To better understand this variation at the level of individual genes in each cell type sample, we performed a differential gene expression (DGE) analysis of *hpo* and *yki* knockdown conditions compared to the corresponding control samples at each stage. We excluded early-stage germ line *hpo* and *yki* RNAi samples from this analysis, because we did not recover enough cells from these stages to make robust conclusions about gene expression levels. In general, we found more differentially expressed genes in germ cells than in somatic cells, when either *hpo* or *yki* levels were reduced (Figure 2D, Supplemental Table S3).

In the soma, we detected 17.7% more upregulated (365) than downregulated (310) genes under *hpo* knockdown, compared to 61.5% more upregulated (231) than downregulated (143) genes under *yki* knockdown (Figure 2D, Supplemental Table S3). In contrast, in the germ cells we observed on average over twice as many down-regulated genes (1,161 and 1,120 under *hpo* and *yki* knockdown, respectively) as upregulated genes (421 and 600 under *hpo* and *yki* RNAi, respectively) in both knockdown conditions. Interestingly, this much larger number of differentially expressed *hpo*-and *yki*-responsive genes in the germ line did not correspond with a notable change of the germ line transcriptional landscape throughout the developmental period under study (Figure 2C). The soma, on the other hand, exhibited distinct gene expression responses to Hpo signaling over developmental time (Figure 2B), and these differential responses were explained by a comparatively small number of genes (Figure 2D). We had previously observed that in the somatic cells, the cell proliferation phenotypes induced by the reduction of *hpo* and *yki* activity were diametrically opposed (Sarikaya and Extavour, 2015). Consistently, differential gene expression analysis of *hpo* and *yki* knockdown conditions showed that in the soma, most of the differentially regulated genes (75%) were exclusive to each genotype (*hpo RNAi* 55% and *yki* RNAi 20%) (Figure 2E). Of all the differentially expressed genes, the expression of only four genes (*apterous*, *CG11400*, *wrapper*, and *expansion*) was significantly affected in opposite ways by *hpo* and *yki* RNAi treatment (Figure 2E). By contrast, in the germ cells the majority (51%) of the differentially expressed genes were common to both *hpo* and *yki* loss of function conditions (869 commonly downregulated and 245 commonly upregulated), and only one gene (*CG34408*) showed an “opposite” expression change (upregulated in *yki* loss of function and downregulated in *hpo* loss of function) (Figure 2F). These results are consistent with the above-described observations that *yki* and *hpo* knockdowns induced similar changes in the germ line transcriptome (Figure 2C), but induced genotype-specific effects on the transcriptome of the soma (Figure 2B).

### Functional prediction analysis of the differentially expressed genes

The differentially expressed genes in the germ cells included more significantly enriched pathways from the Kyoto Encyclopedia of Genes and Genomes (KEGG) and terms from the Gene Ontology (GO) database than the genes differentially expressed in the soma (Figure 3, Supplemental Figure S2). We found that the Hippo, MAPK, and Hedgehog pathways were significantly enriched among the downregulated genes in the germ line under both *hpo* and *yki* knockdown conditions (Figure 3A). We additionally found that DNA replication and mismatch repair pathways were enriched within the genes upregulated in response to *hpo* knockdown in germ cells (Figure 3A). In the germ line response to *yki* knockdown, we observed enriched GO terms relevant to meiosis and cell cycle regulation, particularly in late stage ovaries (Figure 3C, Supplemental Figure S2).

**Figure 3:**
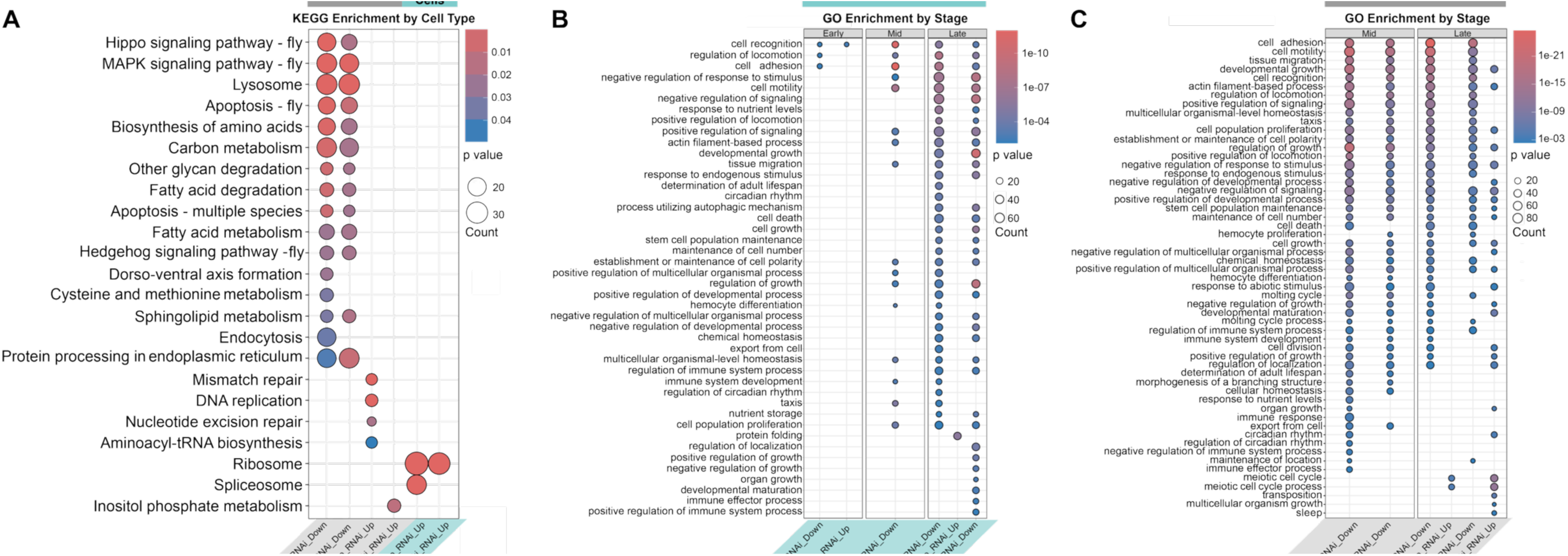
Perturbing *hpo* or *yki* levels has a larger transcriptomic impact on germ cells than somatic cells. **(A)** Significantly enriched (Benjamini-Hochberg adjusted p-value<0.05) KEGG pathways within the groups of significantly upregulated (Up) and downregulated (Down) genes in germ cells and somatic cells under *hpo* and *yki* RNAi conditions (grouping mid and late stages together). **(B, C)** Significantly enriched (Benjamini-Hochberg adjusted p-value < 0.01 and q-value < 0.01) GO-terms of Biological Process level 3 within the significantly upregulated (Up) and downregulated (Down) genes in somatic cells (B) and germ cells (C) under *hpo* and *yki* RNAi.

In contrast, somatic ovarian cells displayed very few significantly altered pathways under either knockdown condition (Figure 3A). This is further corroborated by GO analysis and particularly evident in early and mid stage ovaries (Figure 3B). Moreover, GO analysis revealed that members of some of the same processes that were downregulated in the germ line, were also downregulated in somatic *hpo* knockdowns, but with fewer gene counts than those in germ cells. This was the case even in late stage somatic cells, which had the highest number of enriched processes for somatic cells (Figure 3B, Supplemental Figure S2). These processes included cell adhesion, taxis, motility, tissue growth, cell proliferation, stem cell population maintenance, maintenance of cell number, cell proliferation, and cell death (Figure 3B).

### *yki* is downregulated under *hpo* knockdown in the germ line but not the soma

Our previous work suggested that Yki functions downstream of Hpo in the soma but independently of Hpo in the germ line (Sarikaya and Extavour, 2015). We used our cell-type-specific transcriptomes to determine whether this functional difference was reflected at the level of transcript regulation in one or both of germ line and soma. As expected, we observed a significant reduction (adjusted p-value<0.05) of *hpo* transcript levels across all stages and in both cell types under *hpo* knockdown (Figure 4B). The levels of *yki* transcript were also down-regulated (adjusted p-value<0.05) by *yki* RNAi in all stages and in both cell types, with two apparent exceptions (Figure 4C). First, we noted that the 1.4-fold downregulation of *yki* under *yki* RNAi in early-stage somatic cells just missed our significance threshold (adjusted p-value 0.055). Furthermore, somatic cells at this stage displayed a different transcriptional landscape than controls, consistent with *yki* knockdown (Figure 2B). Second, we noted that one of the control samples for mid stage germ cells had anomalously low levels of *yki* transcript compared to other control samples of the same and other stages (outlined in red in Figure 4C). Removal of this putative outlier revealed a significant reduction of *yki* under *yki* RNAi (Supplemental Figure S3B). Thus, we conclude that both RNAi conditions were able to significantly reduce the transcript levels of their respective target genes, and that our analytical methods were able to detect this reduction.

**Figure 4:**
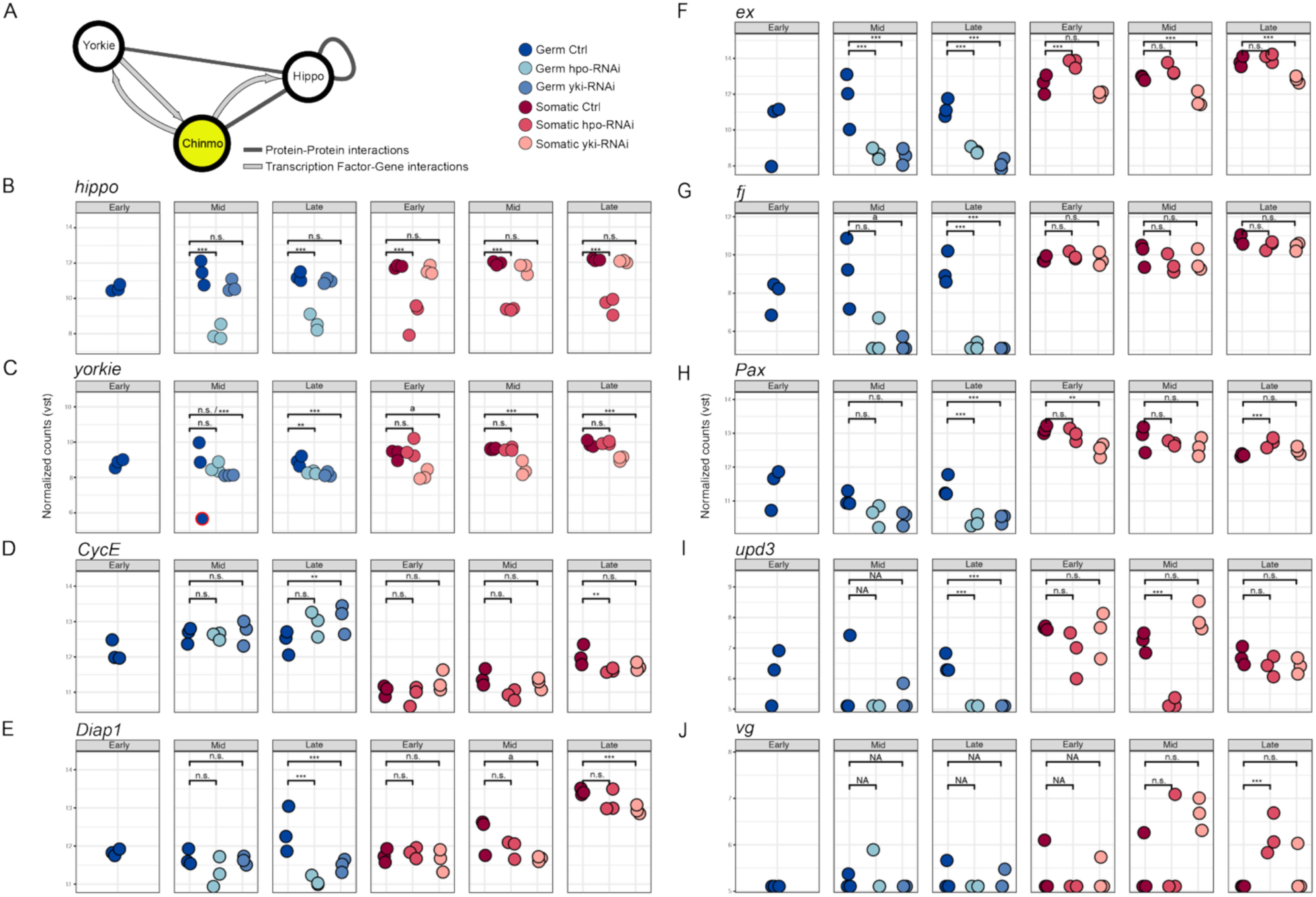
Prediction of *chinmo* as a novel germ cell-specific candidate regulator of *hpo* and/or *yki*. **(A)** Graph showing *hpo*, *yki*, and their only common direct neighbor in our network of protein-protein interactions, *chinmo*. Protein-protein candidate interaction edges are represented by dark grey lines. Transcription factor-gene candidate interactions are represented by light grey arrows pointing to the gene predicted to be regulated by the transcription factor. The transcription factor Chinmo has the potential to transcriptionally regulate both *hpo* and *yki*, can physically interact with Hpo, and can potentially be regulated transcriptionally by Yki. Expression of **(B)** *hpo* and **(C)** *yki* as vst-normalized counts in germ cells (left panels, blue dots) and somatic cells (right panels, red dots) in each RNA-Seq library (color shades indicate genetic condition). The *hpo* and *yki* transcripts appear significantly downregulated in their corresponding RNAi conditions. The *yki* reduction in germ cells mid stages is significant when the control sample with anomalously low levels of *yki* (circled in red) is removed (Supplemental Figure S3). Additionally, *yki* appears significantly downregulated under *hpo* knockdown. Expression of canonical *yki* targets **(D)** *CycE,* **(E)** *Diap1*, **(F)** *ex*, **(G)** *fj*, **(H)** *Pax*, **(I)** *upd3*, and **(J)** *vg* as vst-normalized counts in germ cells (left panels, blue dots) and somatic cells (right panels, red dots) in each RNA-Seq library (color shades indicate genetic condition). Statistical significance levels from the differential expression analysis are codified as *** = adjusted p-value<0.01; ** = adjusted p-value<0.05; a = adjusted p-value = 0.055, n.s. = not significant, NA = values did not pass DESeq2 filtering.

In terms of the effect of RNAi against *yki* or *hpo* on each other’s transcript levels, we found that *yki* transcript levels were significantly reduced under *hpo* RNAi in late stage germ cells, but not at earlier stages, and not in any stage in the soma (Figure 4C). *yki* RNAi (log2FC=-1) and the *hpo* RNAi (log2FC=-0.8) conditions caused a similar reduction in *yki* transcript in the germ line at this late stage. In contrast, the knockdown of *yki* did not yield a significant change in *hpo* transcript levels in either cell type at any stage (Figure 4B). A possible exception may be the mid stage germ cells: when the control sample with anomalously low levels of *yki* is excluded, *hpo* appears significantly downregulated under *yki* knockdown with a fold change of -1.1 (Supplemental Figure S3B). The degree of possible down-regulation of *hpo* in the *yki* RNAi condition is much weaker (Log2FC=-1.1) than the reduction of *hpo* produced by the *hpo* RNAi condition at the same stage (Log2FC=-4.1) (Figure 4B). Taken together, our results suggest that the transcriptional regulatory relationship between *hpo* and *yki* in the larval female germ line is complex, different from that in the soma, developmental stage-specific, and not directly predicted from previous observations on the regulatory interactions of their protein products.

We examined the relative expression levels of seven canonical, positively regulated Yki target genes: *Cyclin E* (*CycE*), *Death-associated inhibitor of apoptosis 1* (*Diap1*), *expanded* (*ex*), *four-jointed* (*fj*), *Paxillin* (*Pax*), *unpaired 3* (*upd3*), and *vestigial* (*vg*), in each RNA-Seq library. All of the examined genes exhibited significant changes in transcript levels in at least one condition. Of these, all but *CycE* were downregulated under *yki* knockdown, consistent with the canonical understanding of the Hpo pathway (Figure 4E-J). Under *hpo* knockdown, however, germ cells and somatic cells exhibited different transcriptional responses. Of genes differentially expressed in the soma, all but *CycE* and *upd3* were upregulated in the *hpo* knockdown condition (Figure 4D-J). In germ cells, all target genes except for *CycE* were downregulated under *hpo* knockdown, most consistently in late stage ovaries (Figure 4D-J). This downregulation is inconsistent with the canonical model of the Hpo pathway, suggesting a possible Yki-independent regulation of Yki target genes by Hpo and a more complex interaction network between Hpo pathway components than previously expected. Notably, in the examined *yki* target genes, *hpo* and *yki* knockdowns induced similar effects in the germ line, but contrasting effects in the soma (Figure 4D-J).

### Gene regulatory network analysis suggests *chinmo* as a novel mediator of *hpo*-dependent *yki* regulation

To generate hypotheses as to how *hpo* might regulate *yki* expression in the larval germ line, we performed a network analysis. To this end, we built a directed network using the DroID database of protein-protein interactions and transcription factor-encoding genes (Murali et al., 2011; Yu et al., 2008). Additionally, we used publicly available Yki and GAF Chip-seq data from *D. melanogaster* embryo, wing disc and S2 cells (Oh et al., 2013) to find possible direct targets of Yki in *D. melanogaster* and added them into the network. From the ChIP-seq Yki data, we identified 1,513 peaks of Yki-binding regions, which we mapped to the closest gene, thus identifying 1,118 genes putatively regulated by Yki across the three studied cell types. By taking the intersection of sets of putative Yki targets from all three cell types, we hoped to retrieve targets of Yki that might also be relevant in the ovary.

In the germ line, 31.4% and 33.4% respectively of all the genes downregulated in either *hpo* or *yki* knockdowns were putative direct targets of Yki (Supplemental Figure S4A). Conversely, of the 1,118 putative Yki targets in our network that were differentially expressed, most of them were downregulated (94.59% under *hpo* RNAi and 79.14% under *yki* RNAi) (Supplemental Figure S4B). In contrast, in somatic cells only 8.23% and 2.86% of putative Yki target genes were significantly affected by *hpo* and *yki* knockdowns respectively (Supplemental Figure S4A). Similarly, among the differentially expressed putative Yki targets the downregulated fraction was much lower in the somatic cells than observed in the germ cells (67.37% were downregulated under *hpo* knockdown, and 50% under *yki* knockdown; Supplemental Figure S4B). This is consistent with the overall much higher number of differentially regulated *hpo*- and *yki*-responsive genes in the germ cells relative to somatic cells (Figure 2D).

To investigate the potential regulatory landscape associated with *hpo* and *yki* in the developing ovary, we examined their surrounding protein–protein interaction network. This analysis revealed several interactors with established roles in developmental regulation, particularly in transcriptional regulation and chromatin state modulation. Among these were the protein products of the patterning genes *caudal* and *dorsal*, as well as chromatin-associated factors including Trithorax-like, BEAF-32, and Cp190 (bric-a-brac domain binding). Although these proteins were closely associated with the Hpo–Yki network, they were connected indirectly as second or third-degree nodal interactors. We found that the shortest path linking *hpo* and *yki* in the network was through a predicted interaction of Hpo with a transcription factor called Chronologically inappropriate morphogenesis (Chinmo) (Zhu et al., 2006), which was predicted to regulate *yki* expression (Figure 4A). Moreover, Chinmo was the sole candidate in our interaction network that was predicted to directly link both Hpo and Yki (Figure 4A). Our RNA-Seq data showed that this transcription factor was significantly upregulated in late stage germ cells under both *hpo* and *yki* loss of function conditions, and conversely, was significantly downregulated in late stage somatic cells under both RNAi conditions (Supplemental Figure S5). We therefore considered the hypothesis that differential *chinmo* expression and/or function between germ line and soma might contribute to the different responses of those cell types to Hpo pathway activity and sought to test this hypothesis experimentally.

### Nuclear Chinmo in the Germ cells is regulated by *yki* expression

Our RNA-Seq results indicated that *chinmo* is expressed at constant levels in wild type germ line cells throughout the developmental stages studied here, while levels of *chinmo* decreased over developmental time in the ovarian somatic cells (Supplemental Figure S5). Given that *chinmo* encodes a transcription factor (Chen et al., 2020; Zhu et al., 2006), we used an anti-Chinmo antibody to examine the expression of Chinmo protein in the cells of the larval ovary. We detected high levels of Chinmo protein in the nuclei of the germ cells in mid and late stages of larval development, but only weak expression in the somatic ovarian cells (Figure 5A; Supplemental Figure S6).

**Figure 5.**
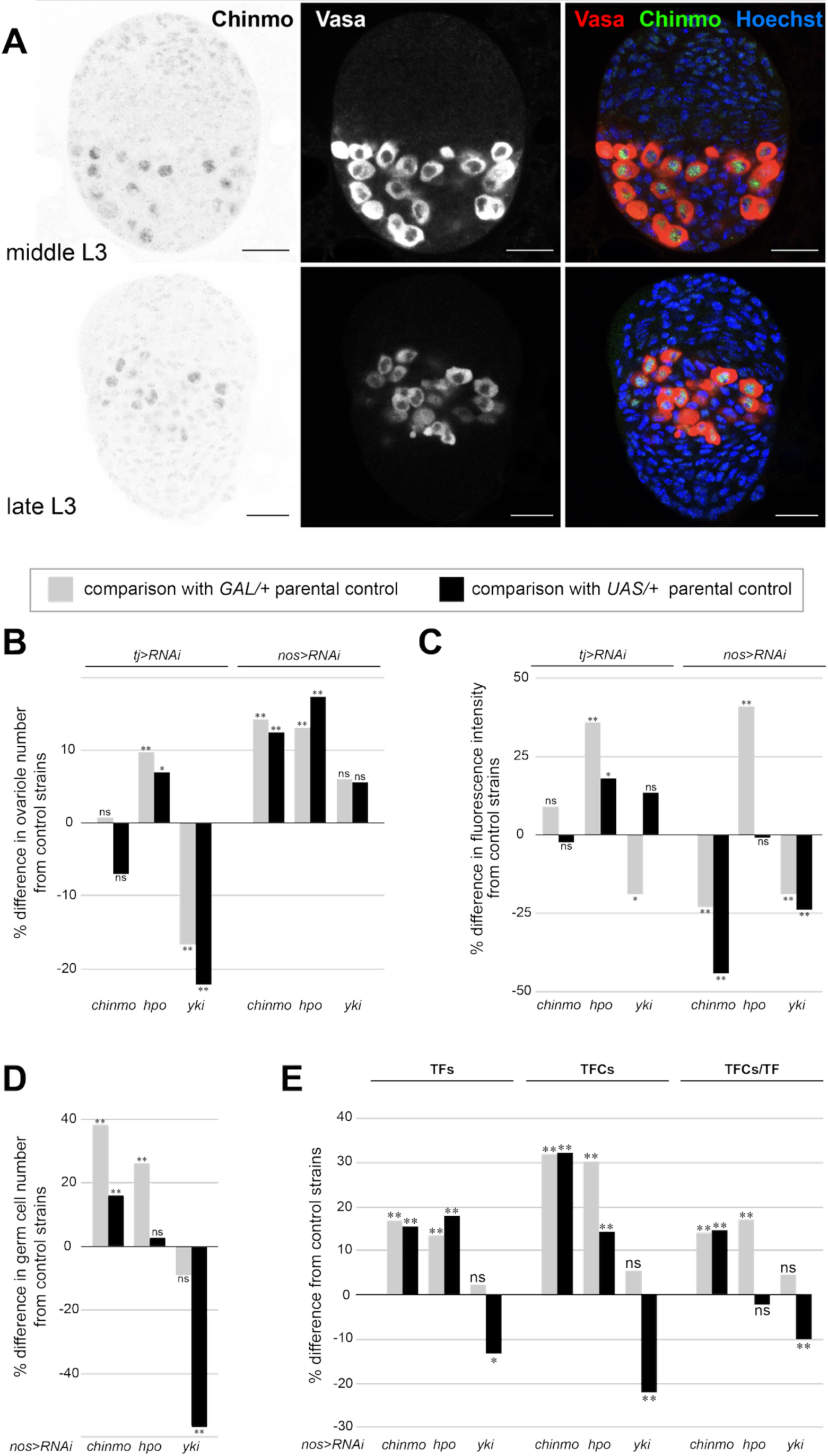
Chinmo regulates germ cell number autonomously and somatic cell number and morphogenesis non-autonomously in the developing ovary. **(A)** Chinmo is detected in germ cell nuclei at mid and late stages of larval development. Red = Vasa; Chinmo = green, scale bars = 20μm. **(B)** *chinmo RNAi* in the germ cells causes an increase in ovariole number. Percentage difference in adult ovariole number where *UAS:chinmo[RNAi]*, *UAS:hpo[RNAi],* or *UAS:yki [RNAi]* are driven in the somatic cells (by *tj:GAL4*) or in the germ cells (by *nos:GAL4*). n=20 for each genotype. **(C)** Abrogation of Hpo signaling in somatic cells causes a change in Chinmo levels in germ cell nuclei. Percentage difference in detected fluorescence levels of Chinmo protein in larval ovarian germ cell nuclei when *UAS:chinmo[RNAi]*, *UAS:hpo[RNAi]* or *UAS:yki[RNAi]* are driven in the somatic cells or in the germ cells. N ≥ 50 nuclei from five ovaries. **(D)** *chinmo[RNAi]* in germ cells causes an increase in germ cell number. Percentage difference in germ cell number where *UAS:chinmo[RNAi]*, *UAS:hpo[RNAi]* or *UAS:yki[RNAi]* are driven in germ cells. **(E)** chinmo[RNAi] in germ cells causes an increase in Terminal Filament (TF) number, Total Terminal Filament Cell (TFC) number and in the number of TFCs per TF. Percentage differences in TF number and TFC number in each ovary, and the number of TFCs per TF when *UAS:chinmo[RNAi]*, *UAS:hpo[RNAi]* or *UAS:yki[RNAi]* were driven in the germ cells. In **B** through **E**, grey bars = comparisons to *GAL4/+* parental controls; blacks bars = comparisons to *UAS/+* parental controls. Statistical significance was calculated using a student’s two-tailed t-test with unequal variance. (** = p<0.01, * = p<0.05, ns = p>0.05).

To determine the functional significance of *chinmo* expression in the larval ovary, we reduced *chinmo* expression in either the somatic cells or the germ line cells using RNAi knockdowns. We used ovariole number as a measure of effective change in phenotype in comparison with the *RNAi* knockdowns of our predicted interactors of Chinmo in this context, Hpo and Yki, both of which we previously identified as regulating ovariole number (Sarikaya and Extavour, 2015). We found that, consistent with the very low levels of detectable Chinmo protein in the soma, reduction of *chinmo* in the somatic cells, unlike the reduction of either *hpo* or *yki,* caused no significant change in ovariole number (Figure 5B; Supplemental Table S4). In contrast, abrogation of *chinmo* in the germ cells caused a significant increase in ovariole number, as did reduction of *hpo* in the germ cells (Figure 5B; Supplemental Table S4). Reducing levels of y*ki* in the germ cells did not cause any change in ovariole number (Figure 5B; Supplemental Table S4).

Our network model predicted a putative transcriptional regulatory relationship between *yki* and *chinmo*, and a potential protein-protein interaction between Hpo and Chinmo proteins. We therefore asked whether levels of *yki* or *hpo* gene activity influenced the levels of Chinmo protein in the nuclei of germ cells in the developing ovary. Consistent with the methods used in our previous studies of ovariole number (Sarikaya et al., 2019, 2012; Sarikaya and Extavour, 2015), in this case and in subsequent quantifications, we report a result as statistically significant only when it differs significantly from both parental controls (*GAL4* and *RNAi* parental lines). Although conditions where comparisons to only one control is statistically significant can reveal differences from just one of the parental backgrounds, we do not use them here as statistically supported evidence to draw our conclusions. We chose this approach because ovariole number is a highly polygenic trait that is strongly influenced by genetic background (e.g. Lobell et al., 2017; Wayne et al., 2001). Thus, for the purposes of determining potential gene regulatory relationships relevant to ovariole number determination, here we wished to focus only on those changes that were statistically significantly different from both parental control genetic backgrounds.

Consistent with our network model, Chinmo protein levels were not significantly affected by *hpo RNAi* in germ cells when compared to both parental controls (Figure 5C; Supplemental Figure S6A). Similarly, reduction of *chinmo* or *yki* expression in somatic cells did not significantly alter detectable Chinmo protein levels in germ cells (Figure 5C; Supplemental Table S4; Supplemental Figure S6B-C). However, Chinmo protein levels did significantly increase in the germ cells upon *hpo* knockdown in somatic cells (Figure 5C; Supplemental Table S4; Supplemental Figure S6A), indicating a non-cell autonomous role for Hpo in Chinmo regulation in the germ line. Reduction of *chinmo* or *yki* in the germ line caused a reduction in detectable Chinmo protein in these cells (Figure 5C; Supplemental Table S4; Supplemental Figure S6B-C).

To understand the functional impact on the ovary of Chinmo levels in the larval germ cells (Figure 5C; Supplemental Table S4), we quantified the number of germ cells, TFCs, and TFs at the late larval stage, and ovarioles in the adult, under loss of germ line expression of *chinmo*, *yki* and *hpo*. Reduction of *chinmo*, but not *yki* or *hpo*, in the germ cells led to a significant increase in germ cell number when compared to both *GAL4* and *UAS* controls (Figure 5D; Supplemental Table S4; Supplemental Figure S6D). We found that the increase in ovariole number caused by reduction of *chinmo* or *hpo* in the germ cells (Figure 5B; Supplemental Table S4) was the result of a non-cell-autonomous increase in TFC number and in TF number (Figure 5E; Supplemental Table S4). Interestingly, reduction of *chinmo* levels in the germ line also caused a significant increase in the number of TFCs in each TF (Figure 5E; Supplemental Table S4). Consistent with the observation that *yki* reduction in germ cells did not change ovariole number (Figure 5B; Supplemental Table S4), this genetic background also had wild type numbers of TFCs and TFs (Figure 5E; Supplemental Table S4).

Taken together, these data are consistent with the hypothesis that Chinmo protein in the nucleus of germ cells regulates germ cell number cell-autonomously, and that Chinmo protein levels in the germ line require germ line expression of *yki*. Further, expression of *chinmo* and *hippo* in the germ cells also exerts a non-cell autonomous effect on somatic cells, limiting TFC number, TF number, and hence ovariole number.

## DISCUSSION

We have provided evidence for a novel genetic mechanistic hypothesis to explain the differential response of germ line and somatic cells to the Hpo signaling pathway during the larval development of the *D. melanogaster* ovary. We propose that the transcription factor Chinmo acts as a modifier of Hpo-independent cell number regulation of the germ line, but not of the somatic gonad cells. Future experiments will be needed to elucidate the molecular mechanisms through which Chinmo interacts with various members of the Hpo pathway.

### The Hippo pathway has distinct effects on transcription in the germ cells and the soma of the larval ovary

In this study, we knocked down either *hpo* or *yki* using RNAi driven in either the germ cells or the somatic cells, thus providing us with a lens into the effects of each of these genetic manipulations on the transcriptomic landscapes in each cell type. We chose the specific RNAi reagents used in this study for consistency with our previous study, in which we showed that they had the same relevant phenotypes as the respective null alleles (Sarikaya and Extavour, 2015). However, it is worth noting that these *yki* and *hpo* RNAi conditions do not completely eliminate detectable transcripts of the target genes. Additionally, we have not performed experiments to rule out off-target effects of our knockdowns.

The canonical understanding of the Hippo signaling pathway suggests that the presence of phosphorylated Hpo in the cellular milieu prevents the nuclear localization of Yki and therefore the downstream expression of its target genes. Under this model, the absence of Hpo in the cytoplasm allows Yki to remain unphosphorylated, enter the nucleus, and drive the expression of its downstream targets. This model therefore predicts a different transcriptional profile with and without Hpo activity. Our data indicate that knockdown of either *hpo* or *yki* in the soma impacts transcript levels of two mutually exclusive sets of genes (Figure 2B). In the germ cells, however, this is not the case: the effects of the knockdown of either *hpo* or *yki* caused transcriptional changes in a similar set of genes (Figure 2C). This suggests that *hpo* and *yki* RNAi have different effects on the transcriptome of the soma, but similar effects on germ line. This interpretation is consistent with the analysis of canonical Yki target expression in the knockdown conditions, wherein most targets had contrasting effects in the soma, but similar effects in the germ line, with a few notable exceptions (Figure 4D-J). Specifically, *CycE* was upregulated under *yki* RNAi in late stage germ cells, and downregulated under *hpo* RNAi in late stage somatic cells, contrasting with expectations of the Hpo canonical pathway (Figure 4D). Similarly, *upd3* was downregulated under *hpo* RNAi in mid stage somatic cells.

Furthermore, we found that more genes were differentially expressed in the germ line than in the soma (Figure 2D). The higher number of significantly upregulated *yki*-regulated genes in somatic tissues could mean that *yki* directly or indirectly represses more genes in somatic tissue than it activates. Again, we observed differences in the effect of *hpo* and *yki* knockdown on germ cells and somatic cells. The higher number of downregulated genes in the germ cells under these conditions suggests that *hpo* and *yki* may operate in germ line tissue primarily through promoting the transcription of genes rather than repressing them. In all genetic backgrounds, we observed that most of the transcriptional variation in the soma was attributable to the progression of development (Figure 2B), while in the germ line, the greatest variation between the samples was due to abrogation of Hippo signaling pathway function (Figure 2C). This suggests that many of the transcriptome differences we observed in the soma across development reflect the dynamic morphogenetic changes taking place in the somatic cells of the ovary during the process of terminal filament determination and organization.

### *Hpo* RNAi in germ cells does not affect cell number via *yki* repression

The similar transcriptomic impact of *yki* and *hpo* knockdowns in the germ cells is intriguing given their opposite phenotypic effects on cell proliferation in the soma (Sarikaya and Extavour, 2015). We had previously shown that *hpo* knockdown in somatic cells increased their number, while *yki* knockdown decreased their number (Sarikaya and Extavour, 2015). Similarly, overexpression of *hpo* and *yki* in somatic cells and in germ cells also had opposite phenotypic effects, whereby *hpo* overexpression reduced somatic cell number and *yki* knockdown increased it (Sarikaya and Extavour, 2015). These results were consistent with the canonical model of Hippo pathway activity. However, although *yki* knockdown in germ cells decreased germ cell number, *hpo* knockdown did not change it (Sarikaya and Extavour, 2015). We suggest that transcriptional regulation of *yki* by *hpo* may contribute to the previously reported apparent imperviousness of germ cell number to *hpo* knockdown. We noted that in the germ line, a reduction in *hpo* caused a reduction in *yki* (Figure 4C), but there was no detectable regulation of *hpo* by *yki* (Figure 4B). To our knowledge, this is the first reported example of the transcriptional regulation of *yki* by *hpo.* We hypothesize that when *hpo* is reduced in germ cells, although nuclear localization of Yki is not inhibited, there is less *yki* transcript. This could in principle lead to less Yki protein and less efficient triggering of the transcriptional cascade that would result in a cell number increase. In sum, we hypothesize that a balance of *hpo* and *yki* transcript levels contributes to regulating the number of germ cells.

### *Chinmo* and its role in the regulation of *hpo* and *yki*

Chinmo is a BTB-Zinc finger transcription factor that is conserved across insects (Nadimpalli et al., 2015; Spokony and Restifo, 2007). Chinmo was first identified as a negative regulator of the JAK/STAT signalling pathway (Fisher et al., 2012; Flaherty et al., 2009; Wang et al., 2013) and acts as a transcriptional repressor downregulating the expression of a broad spectrum of genes including *shotgun* (*E-Cadherin*) (Enomoto et al., 2021), *β Spectrin*, *polychaetoid*, *scribble* and *mirror* (Grmai et al., 2021). In some studied contexts, Chinmo prevents the differentiation of cells and promotes their ability to proliferate(Chafino et al., 2023).

*chinmo* is the only gene we found in our network analysis that was predicted to directly regulate *yki* in response to *hpo* levels, since Hpo and Chinmo proteins have the potential to interact based on yeast two hybrid data (Yu et al., 2008). However, these potential interactions will require experimental validation in future studies to confirm their presence in the context of the developing ovary. Furthermore, our network predicts that *chinmo* has the potential to regulate the expression of both *hpo* and *yki* (Figure 4A), and that in turn, *yki* could regulate the expression of Chinmo based on ChIP-seq data (Oh et al., 2013). We hypothesize that *chinmo* could be involved in regulating *yki* transcript levels in response to Hpo activity levels, which would provide a mode of control of Hpo by Yki through Chinmo. Previous studies of *chinmo* in the *D. melanogaster* testis and ovaries showed that it plays a key role in inducing male fate in somatic cells of the testis, but that it is not required in ovarian somatic cells (Ma et al., 2016). Consistent with that report, our results show increased expression of *chinmo* in response to *yki* knockdown in the germ line but not in the soma (Supplemental Figure S5).

Given the low levels of detectable Chinmo in somatic nuclei of the larval ovary (Figure 5A), and that adult ovariole number is unchanged by somatic reduction of *chinmo* (Figure 5B), we hypothesize that Chinmo does not act in somatic cells to impact the known regulation of the number of TFCs, TFs or ovarioles by *hpo* or *yki* (Sarikaya and Extavour, 2015). In contrast, we detected Chinmo protein in larval germ cell nuclei (Figure 5A), where its expression appears to be dependent on *yki* levels in the germ cells (Figure 5C). Further, we observed that reduction of *chinmo* in germ cells caused an increase in germ cell number (Figure 5D), suggesting that Chinmo acts in larval ovarian germ cells to limit their proliferation. This is reminiscent of the observation that in *chinmo^ST^* loss of function mutants, adult male germ cells overproliferate (Ma et al., 2014). On the other hand, *chinmo^-/-^* clones in a heterozygous *chinmo^+/-^* background do not overproliferate (Tseng et al., 2021). Thus, the cell-autonomous impact of *chinmo* function on germ cell proliferation appears to be context- and sex-specific. Combining the results presented herein with our previous understanding of Hippo signaling in the ovary, we propose a model for the cell type-specific regulation of *hpo*-mediated proliferation during larval ovary development (Figure 6).

**Figure 6.**
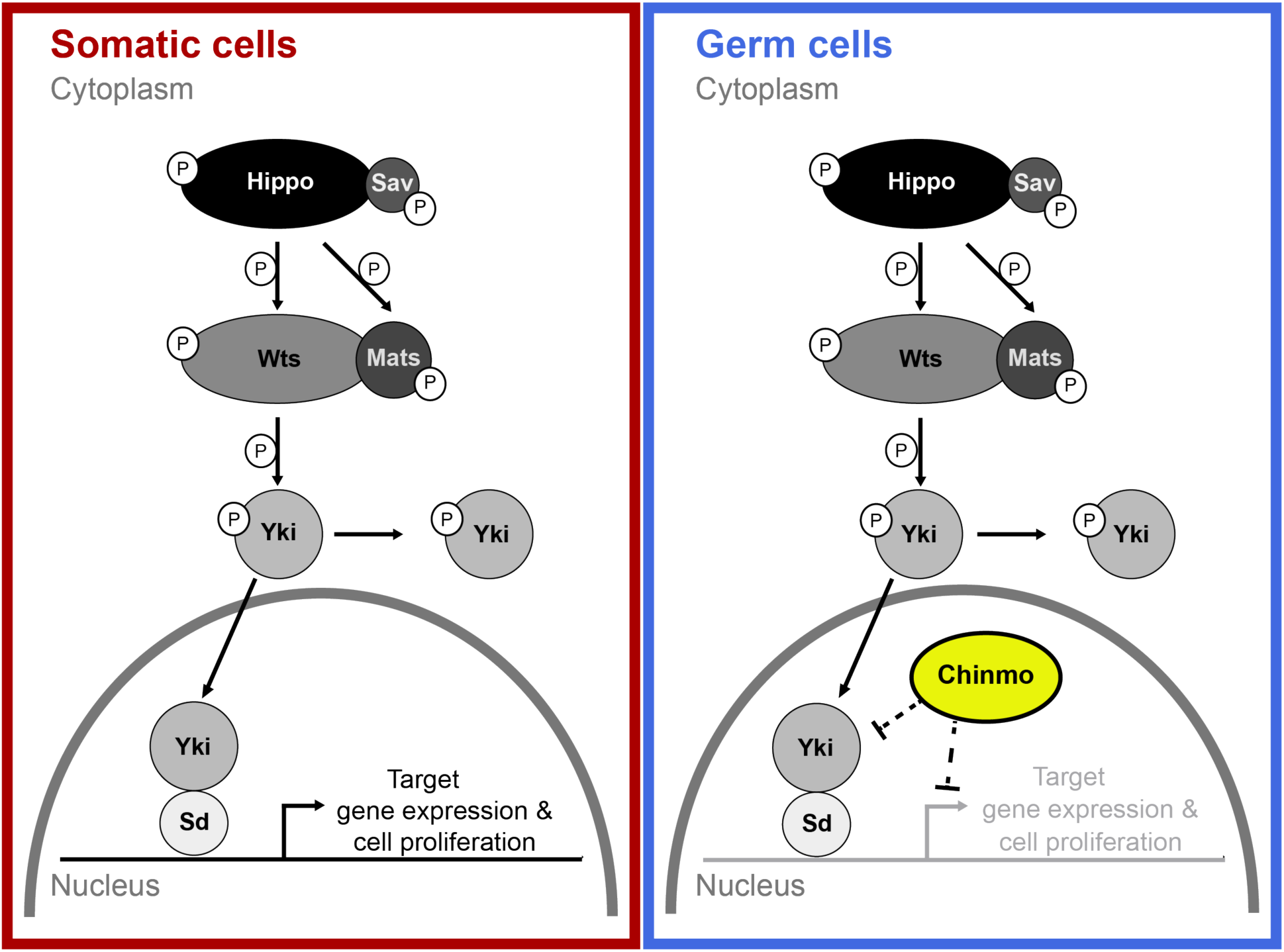
Hypothesis: Chinmo regulates Hpo signaling activity in the germ cells but not somatic cells. We previously showed that the components of the Hippo signaling pathway are present in both the germ and somatic cells (Sarikaya and Extavour, 2015), and that the somatic cells (red) function under the influence of the canonical Hippo pathway (Sarikaya and Extavour, 2015). Under this model, the reduction of either *hpo* or *yki* causes a change in germ cell number. In germ cells (blue), Chinmo is present, which adds a secondary Hippo-independent repression of *yki*-dependent cell proliferation. Because our data cannot determine whether Chinmo acts directly on the localization of Yki to the nucleus, or indirectly by repressing cell proliferation downstream of Yki, the Chinmo-Yki relationship is indicated by dashed lines.

In this model, Chinmo and Yki function in opposition to each other in the germ line to regulate germ cell number, limiting or promoting germ cell proliferation respectively. Thus, unlike in the somatic cells, loss of *hpo* in germ cells does not cause an increase in germ cell number because Chinmo continues to prevent Yki-dependent transcription. The similarities that we observed between the transcriptional profiles of the *hpo* and *yki* knockdown in the germ cells might be explained by the prediction that Chinmo prevents Yki-dependent transcription in these cells. Thus, we propose that the interplay of Chinmo and Yki levels in the nucleus contributes to the maintenance of germ cell number, and that disrupting this balance causes a change in germ cell number. The model proposed in this study is also consistent with the Hpo-independent function of Yki in the germ cells that was suggested by our previous work (Sarikaya and Extavour, 2015). However, the precise molecular mechanism by which Chinmo might inhibit Yki-dependent transcription remains unknown and will require future studies to elucidate. To further understand this connection, future work should include an analysis of canonical Hpo and Yki targets in the context of *chinmo* reduction. Additionally, an analysis of *chinmo* overexpression on the transcriptional landscape of the ovary in a *hpo* or *yki* knockdown background could be informative to assess the sufficiency of Chinmo in modulating cell responses to Hippo signaling.

### Cell-non-autonomous effects of Chinmo and Hippo

Reduction of somatic *hpo* causes a non-autonomous increase in the levels of detectable nuclear Chinmo in the germ cells (Figure 5C). This could be due to *hpo* downregulation in either or both of TFCs or intermingled cells, given that the GAL4 driver we used here (*tj:Gal4*) is expressed in both somatic cell types (Sarikaya and Extavour, 2015). We note that downregulation of *hpo* in intermingled cells causes JAK/STAT pathway upregulation in germ cells (Sarikaya and Extavour, 2015), and that Chinmo is positively regulated by the JAK/STAT signalling pathway (Fisher et al., 2012; Flaherty et al., 2009; Wang et al., 2013). It is thus possible that abrogating *hpo* activity in intermingled cells causes nuclear Chinmo levels to rise in the germ line because of the non-autonomous upregulation of JAK/STAT activity in germ cells. However, the present study does not address the complexities of mechanisms of cell-non-autonomous effects between somatic and germ cells in ovarian development. Future mechanistic investigations of these effects will be necessary to comprehensively address these intercellular dynamics.

Our observation of the non-autonomous impact of germ cell number on multiple aspects of TFC number and organization (Figure 5B, 5E) is a new and intriguing finding. Reducing *chinmo* or *hpo* in the germ line led to a significant increase in TFC, TF (Figure 5E) and ovariole number (Figure 5B), suggesting that germ cell number can positively regulate ovariole number, a hypothesis that has remained underexplored in the field. In our previous work (Sarikaya and Extavour, 2015) we did not observe a statistically significant increase in germ cells in *nos:GAL4>hpo[RNAi]* larval ovaries (Sarikaya and Extavour, 2015), which is the same genetic background we used in the present work. Further, although we did not count ovarioles in our previous analysis of the *nos:GAL4>hpo[RNAi]* condition, we did not observe a statistically significant non-autonomous impact of *hpo* knockdown in the germ line on terminal filament cell number. We attribute these differences to the fact that the control genotypes used for comparison and calculation of statistical significance in the present work (*GAL4* or *UAS* heterozygotes obtained by crossing the parent strains to wild type strain Oregon R), were slightly different from the control genotypes used in our previous study (sibling controls) (Sarikaya and Extavour, 2015). We and others have observed that ovariole number, and its contributing developmental processes, are such highly polygenic traits that even small changes in genetic background can have statistically detectable effects (Bergland et al., 2008; Coyne et al., 1991; Kumar et al., 2020; Orgogozo et al., 2006; Telonis-Scott et al., 2005; Thomas-Orillard, 1976; Wayne et al., 2001; Wayne and McIntyre, 2002).

Although increasing ovariole number by increasing germ cell number has not, to our knowledge, previously been reported, there are at least two reported examples of the inverse situation, where reduced germ cell number is correlated with reduced ovariole number. In one example, ovaries of adult females carrying the dominant female sterile *ovo^D1^* allele, which lacks functional adult female germ cells, have fewer ovarioles than wild type (Extavour, 2000). This could in principle be because of a reduction in total germ cell number during larval stages in this genotype, which is consistent with the interpretation of larval germ line clonal analysis of this allele (Perrimon, 1984). In a second example, agametic adult females generated by irradiating the posterior pole of embryos (Geigy, 1931), have fewer ovarioles than controls (Aboïm, 1945).

Finally, we note that reduction of *chinmo* levels in the germ cells also caused an unexpected increase in the number of TFCs in each TF (Figure 5E). To our knowledge, this is the first reported evidence of changes in germ line gene influencing TFC number and TF morphogenesis. Determining the non-autonomous regulatory mechanism that allows larval germ cells to impact the future fecundity of the adult by influencing somatic morphogenesis of the ovary is an interesting avenue for future research.

## MATERIALS AND METHODS

### Fly stocks

Flies were reared(Sarikaya et al., 2012) at 25°C and 60% humidity on standard *Drosophila* food with yeast in uncrowded vials as previously described (Sarikaya et al., 2012). The “wild type” genetic background referred to herein is Oregon R (Lindsley and Grell, 1968). The following fly lines were obtained from the Bloomington *Drosophila* Stock Center (BDSC): *w[*]; P{w[+mW.hs]=GawB}bab1[Pgal4-2]/TM6B, Tb[1]* (abbreviated herein as *bab:GAL4*; BDSC stock number 6803); *P{w[+mC]=UAS-Dcr-2.D}1, w[1118]; P{w[+mC]=GAL4-nos.NGT}40* (abbreviated herein as *nos:GAL4*; BDSC stock number 25751); *y[1] v [1]; P(y[+t7.7]v[+t1.8]=TRiP. HMS00006) attP2* (abbreviated herein as *hpo[RNAi]*; BDSC stock number 33614) and *y[1] v[1]; P(y[+t7.7] v[+t1.8]=TRiP.HMS00041)attP2* (abbreviated herein as *yki[RNAi]*; BDSC stock number 34067); *y[1] v[1]; P{y[+t7.7] v[+t1.8]=TRiP.JF02341}attP2* (abbreviated herein as *chinmo[RNAi]*; BDSC stock number 26777). *y w; P{w[+mW.hs]=GawB}NP1624* was obtained from the Kyoto Stock Centre (abbreviated herein as *tj:*GAL4; stock number K104–055). *w[1118], P[UAS Stinger]* (abbreviated herein as *UAS:Green Stinger I*, (Barolo et al., 2000) used for GFP expression was a gift from Dr. James Posakony (University of California, San Diego).

Crosses (described below in “Staging larvae”) were set with 100-200 virgin females and 50-100 males in a 180 ml capacity bottle containing50 ml standard Drosophila food with yeast per bottle, one day prior to egg laying. For somatic *chinmo* knockdown via RNAi, we used a *tj:GAL4* driver (Li et al. 2003), rather than the *bab*:*GAL4* driver (Cabrera et al. 2002), as the latter caused lethality when driving *chinmo [RNAi]* (data not shown). A detailed list of experimental crosses and controls used for transcriptome generation can be found in Supplemental Table S5.

### Staging larvae

Uniformly staged larvae were collected from near-synchronously laid eggs. To obtain a desired genotype, crosses were set as described above in “Fly Stocks” at 25°C at 60% humidity and left overnight to mate. Hourly egg collections were set up on 60 mm apple juice-agar plates (9 g agar, 10 g sugar, 100 ml apple juice and 300 ml water) with a pea-sized spread of fresh yeast paste (baker’s yeast granules made into a paste with a drop of tap water). Eggs were collected hourly for eight hours. The first two collection plates were discarded to remove asynchronously laid eggs that may have been retained inside the females following fertilization. Staged first instar larvae were collected into vials 24 hours after egg collection. Larvae at 72h hours After Egg Laying (h AEL) were designated as early stage, at 96h AEL as middle stage and at 120h AEL as late stage of terminal filament development.

### Dissection and dissociation of larval ovaries

Staged larvae were collected for dissection every hour. Fat bodies were dissected out of the larvae using a pair of forceps. Ovaries, located in the center of the length of each fat body, were then dissected free of the fat body using an insulin syringe needle (BD 328418). Ovaries thus dissected clear of fat body were collected in ice-cold DPBS (Thermo Fisher 14190144) and batches of 20-30 ovaries in DPBS were kept on ice for a maximum of four hours until dissociation and subsequent FACS processing. For dissociation, dissected ovaries in ice-cold DPBS were transferred to 0.25% trypsin solution (Thermo Fisher 25200056) for ten minutes at room temperature in the cavity of a glass spot plate (Fisher Scientific 13-748B). They were then transferred to another cavity containing 2.5% liberase (5 g liberase reconstituted in 2 ml nuclease free water; Sigma 5401119001), teased apart with tungsten needles until most of the clumps were separated, and left in the liberase solution (without agitation) at room temperature for ten minutes. The tissue in liberase was pipetted gently to uniformly mix and dissociate the cells. The cell suspension was then transferred to an RNA Lobind tube (Eppendorf 8077-230) and placed on a vortexer for 1 minute. This sample was immediately subjected to fluorescence activated cell sorting (FACS) and the sorted cell suspension was collected in an RNA Lobind tube containing 100-200 µl Trizol (Thermo Fisher 15596026).

### Flow sorting GFP positive cells

The dissociated cell suspension was sorted in a MoFlo Astrios EQ Cell sorter (Beckman Coulter) run with Summit v6.3.1 software. The dissociated cell suspension was diluted and a flow rate of 200 events per second was maintained with < 98% sorting efficiency during the sorting process. A scatter gate (R1) was employed to eliminate debris, and a doublet gate (R2) was used to exclude non-singlet cells. A 488 nm emission Laser was used to excite the GFP, and the collection was at 576 nm. The GFP-positive cells were designated in gate R3 and sorted directly into Trizol (Supplemental Table S1). The resulting cells collected in Trizol were frozen immediately by plunging the tube in liquid nitrogen and then stored at -80°C until RNA extraction. A single replicate consisted of at least 1000 cell counts pooled from FACS runs.

### RNA extraction

Flow sorted cells stored in Trizol at -80°C were thawed at room temperature and lysed with a motorized pellet pestle (Kimble 749540-0000). A Zymo RNA Micro-Prep kit (Zymo Research R2060) was used to isolate RNA from the homogenized Trizol preparations. An equal volume of molecular grade ethanol (Sigma E7023) was added to Trizol and mixed well with a pellet pestle, then pipetted onto a spin column. All centrifugation steps were done at 10,000g for one minute at room temperature. The column was washed with 400 µl Zymo RNA wash buffer and then treated with Zymo DNase (6 U/µl) for 15 minutes at room temperature. The column was then washed twice with 400 µl Zymo RNA Pre-wash buffer and once with Zymo RNA wash-buffer. The RNA was eluted from the column in 55 µl of nuclease-free water (Thermo Fisher 10977015). The RNA obtained was quantified first using a NanoDrop (Model ND1000) spectrophotometer and then using a high sensitivity kit (Thermo Fisher Q32852) on a Qubit 3.0 Fluorometer (Thermo Fisher Q33216). The RNA was also checked for integrity on a high sensitivity tape (Agilent 5067-5579) with an electronic ladder on an Agilent Tapestation 2200 or 4200.

### Library preparation

cDNA libraries were prepared using the Takara Apollo library preparation kit (catalogue # 640096). Extracted RNA samples were checked for quality using Tapestation tapes. 50µl of RNA samples were pipetted into Axygen PCR 8-strip tubes (Fisher Scientific 14-222-252) and processed through PrepX protocols on the Apollo liquid handling system. mRNA was isolated using PrepX PolyA-8 protocol (Takara 640098). The mRNA samples were then processed for cDNA preparation using the PrepX mRNA-8 (Takara 640096) protocol. cDNA products were then amplified for 15 cycles of PCR using longAmp Taq (NEB M0287S). During amplification PrepX RNA-Seq index barcode primers were added to each library to enable multiplexing. The amplified library was then cleaned up using PrepX PCR cleanup-8 protocol with magnetic beads (Aline C-1003). The final cDNA libraries were quantified using a high sensitivity dsDNA kit (Thermo Fisher Q32854) on a Qubit 3.0 Fluorometer (Thermo Fisher Q33216). cDNA content and quality were assessed with D1000 (Agilent 5067-5582) or high sensitivity D1000 tape (Agilent 5067-5584, for low quantities of cDNA) on an Agilent Tapestation 2200 or 4200.

### Sequencing cDNA libraries

Libraries were sequenced on an Illumina HiSeq 2500 sequencer. Single-end 50bp reads were sequenced on a high-throughput flow cell. Libraries of varying concentrations were normalized to be equimolar, the concentrations of which ranged between 2-10nM per lane. All the samples in a flow cell were multiplexed and later separated based on unique prepX indices to yield at least 10 million reads per library. The reads were demultiplexed and trimmed of adapters using the bcl2fastq2 v2.2 pipeline to yield final fastq data files.

Three RNA-Seq biological replicates were obtained for each condition, each of which had at least 10 million reads. The only exception was the early-stage germ line under both *hpo* and *yki* RNAi knockdowns, for which we only obtained a single biological replicate due to the technical challenge of obtaining enough germ line cells from these knockdown ovaries. These single replicates of early germ line stages were not included in the differential expression analysis but are made publicly available as part of the reported dataset (See Data Availability).

### Immunohistochemistry

Larvae were dissected and stained as previously described (Sarikaya et al., 2012; Sarikaya and Extavour, 2015). For all cell counts, samples were dissected at the larval-pupal transition stage. Primary antibodies and concentrations used to stain the larval-pupal transition stage ovaries are as follows: Mouse anti-Engrailed 4D9 (1:40, Developmental Studies Hybridoma Bank (DSHB)), Rabbit anti-Vasa (1:3000, gift from A. Nakamura, Kumamoto University), Rat anti-Chinmo (1:500, gift from E. Bach, New York University) (Wu et al., 2012). Secondary antibodies used were anti-mouse Alexa Fluor 488, anti-rat Alexa Fluor 488, and anti-rabbit Alexa Fluor 568, all used at 1:400, and Hoechst 33342 used at 1:10,000 (stock solution 10 mg/mL). All samples were imaged on a Zeiss LSM 880 inverted laser confocal microscope. For quantification of Chinmo expression levels, all samples were dissected and stained in parallel. Imaging settings, including laser power, detector gain and step sizes, were kept constant for all experimental and appropriate control samples.

### Quantification of ovariole number, cell number and Chinmo protein level

Ovariole number was quantified as previously described (Kumar et al., 2020). Germ cells were counted manually in larval-pupal transition stage ovaries stained for Vasa protein, using the Manual tracking with TrackMate plugin in Fiji (Schindelin et al., 2012; Tinevez et al., 2017). Terminal filaments and terminal filament cells in each terminal filament were counted manually in larval-pupal transition stage ovaries stained for Engrailed, using the Cell counter plugin in Fiji. Total terminal filament cell number per ovary was calculated by adding up the number of terminal filament cells in each terminal filament.

### RNA-Seq data processing

The *D. melanogaster* genome assembly and gene annotations were obtained from FlyBase version dmel_r6.36_FB2020_05 (Larkin et al., 2020). The reads were aligned with RSEM v1.3.3. (Li and Dewey, 2011). Using STAR v2.7.6a as read aligner, (Dobin et al., 2013) we obtained the gene counts in each library. Because some of the tissue-specific biological samples were sequenced in more than one lane or run, the reads were split into multiple fastq files, and the gene counts belonging to the same biological sample were summed. Gene counts in each dataset were normalized with the variance stabilizing transformation (VST) method implemented in the DESeq2 v1.26.0 (Love et al., 2014) R package. Further analyses, such as principal component analysis, hierarchical clustering, and differential expression analysis, were performed in R using the VST-normalized counts.

### Differential expression analysis

The differential expression analyses were performed with DESeq2 v1.26.0 (Love et al., 2014). Genes with a Benjamini-Hochberg adjusted p-value lower than 0.05 were selected as differentially expressed in the corresponding contrast.

### Functional predictions

The Gene Ontology (GO) and Kyoto Encyclopedia of Genes and Genomes (KEGG) pathways enrichment analyses were performed on the differentially expressed genes with the enrichGO and enrichKEGG functions of the clusterProfiler package (v3.14.3) for R (Yu et al., 2012). The GO terms were obtained using the R package AnnotationDbi (Carlson, 2024) with the database org.Dm.eg.db v3.10.0. The GO overrepresentation analysis of biological process was performed against the gene universe of all *D. melanogaster* annotated genes in org.Dm.eg.db, adjusting the p-values with the Benjamini-Hochberg method, adjusted p-value and q-value cutoff of 0.01, and a minimum of 30 genes per term. For the KEGG enrichment analysis, p-values were adjusted by the Benjamini-Hochberg procedure, and an adjusted p-value cutoff of 0.05 was used.

### Analysis of publicly available ChIP-Seq data

We obtained the ChIP-Seq from GEO (GSE46305 and GSE38594) (Oh et al., 2014, 2013). We downloaded the coordinates of the 3,749, 6,491, and 3,991 Yorkie-bound peaks from embryo (E8-16h), wing imaginal discs, and S2 cells respectively. The coordinates of the peaks were based on the *D. melanogaster* genome assembly BDGP release 5, and transformed to coordinates for the latest genome version (release 6) with the lifterOver function. The peaks were annotated with ChIPseeker (v.1.22.1) using *D. melanogaster* genome annotations from FlyBase (v6.36). We defined the promoter region as the 1,500 bp flanking the transcription start site, and 69-75% of the peaks were found within the promoter regions of genes thus defined. In total 2,331, 3,931, and 2,247, genes with Yki-bound sites in the defined promoter region were identified, in embryo, wing imaginal discs, and S2 cells respectively. 4,980 genes were unique (appeared only in one dataset), 1,118 were common across the three datasets, and 2,406 were common to at least two datasets. Following the same methodology, we analyzed the GAF (*Trl*) Chip-Seq peaks (GSE38594) from embryo (E8-16h) and wing imaginal discs. Thus, we associated the 6,449 and 6,302 GAF-bound peaks from embryo and wing imaginal discs respectively to the promoters of 4,086 genes, of which 2,095 were common to both datasets.

### Network analysis

To build a comprehensive network that could generate a hypothesis as to how *hpo* might regulate *yki*, we started by obtaining the Transcription Factor – Gene (TF-gene) interaction network from the Drosophila Interactions database (www.droidb.org, version DroId_v2018_08), containing 157,462 interactions of 12,323 genes. Using igraph (v1.2.6)_for R (Csárdi and Nepusz, 2006), we loaded the database and build a graph with genes as nodes and their interactions as edges. We then added the 1,118 interactions of Yorkie and the 2,095 interactions of GAF predicted from the above ChIP-seq analysis to the graph. After removing redundancies, we had 158,923 edges and 112,562 nodes. Furthermore, we also downloaded 127,353 unique protein-protein interactions (PPI) between 10,632 proteins from DroID v2018_08 and modeled it as a graph. The TF-gene and PPI graphs were joined, yielding a graph with 328,989 edges and 13,337 nodes. In this graph, the edge attributes describe the interaction type (TF-gene or PPI). Those edges representing TF-gene interactions had directionality edges because TFs regulate the target and not vice-versa, while the PPI edges were bi-directional. In this large graph, using igraph’s function “all_shortest_paths,” we obtained all possible shortest paths (taking into consideration the edge directions) in the network through which Hpo could regulate Yki. Chinmo emerged as the only possible connector between *hpo* and *yki*, requiring only two steps.

## Supporting information

SI Tables 1 through 5

## ACKNOWLEDGEMENTS

We thank the flow cytometry and genomics staff at the Bauer Core Facility of Harvard University Faculty of Arts and Sciences for advice on FACS sorting and sequencing, and members of the Extavour lab for helpful discussions. This work was supported by NIH R01 award R01-HD073499-0 to CGE, by funds from Harvard University, and by the Google Cloud Academic Research Credits Program. CGE is a Howard Hughes Medical Institute investigator.

## AUTHOR CONTRIBUTIONS

ST performed all steps in generating individual transcriptome datasets, including experimental design, ovary dissection and dissociation, RNA-Seq library preparation and sequencing. GY performed all statistical and network analyses of transcriptome data. TK performed and analysed genetic experiments testing the role of *chinmo*. AAS contributed to figure creation and revision of results and discussion. CGE conceived of the project, provided oversight and funding. ST, TK, GY and CGE drafted the manuscript. All authors edited the manuscript.

## DATA AVAILABILITY

All the raw data are publicly available at NCBI-Gene Expression Omnibus (GEO) database under the accession code GSE250320. Control datasets for early, mid and late stage germ line and somatic cells are accessible as a part of the NCBI-GEO database under the accession code GSE172015 (GSM5239772-89). The scripts used to process and analyze the data are available at the GitHub repository https://github.com/guillemylla/Ovariole_morphogenesis_RNAseq, commit ID 171064d.

