## Supplemental Information for "Chinmo is a novel regulator of differential response to Hippo signaling within a single developing organ"

Cassandra G. Extavour

Supplemental Material consists of the following:

- Supplemental Figures S1 through S6 (this document)
- Supplemental Table S1 through S5 (legends in this document; tables are separate files):
    - Supplemental Table S1.xlsx
    - Supplemental Table S2.csv
    - Supplemental Table S3.csv
    - Supplemental Table S4.xlsx
    - Supplemental Table S5.xlsx
- Supplemental References

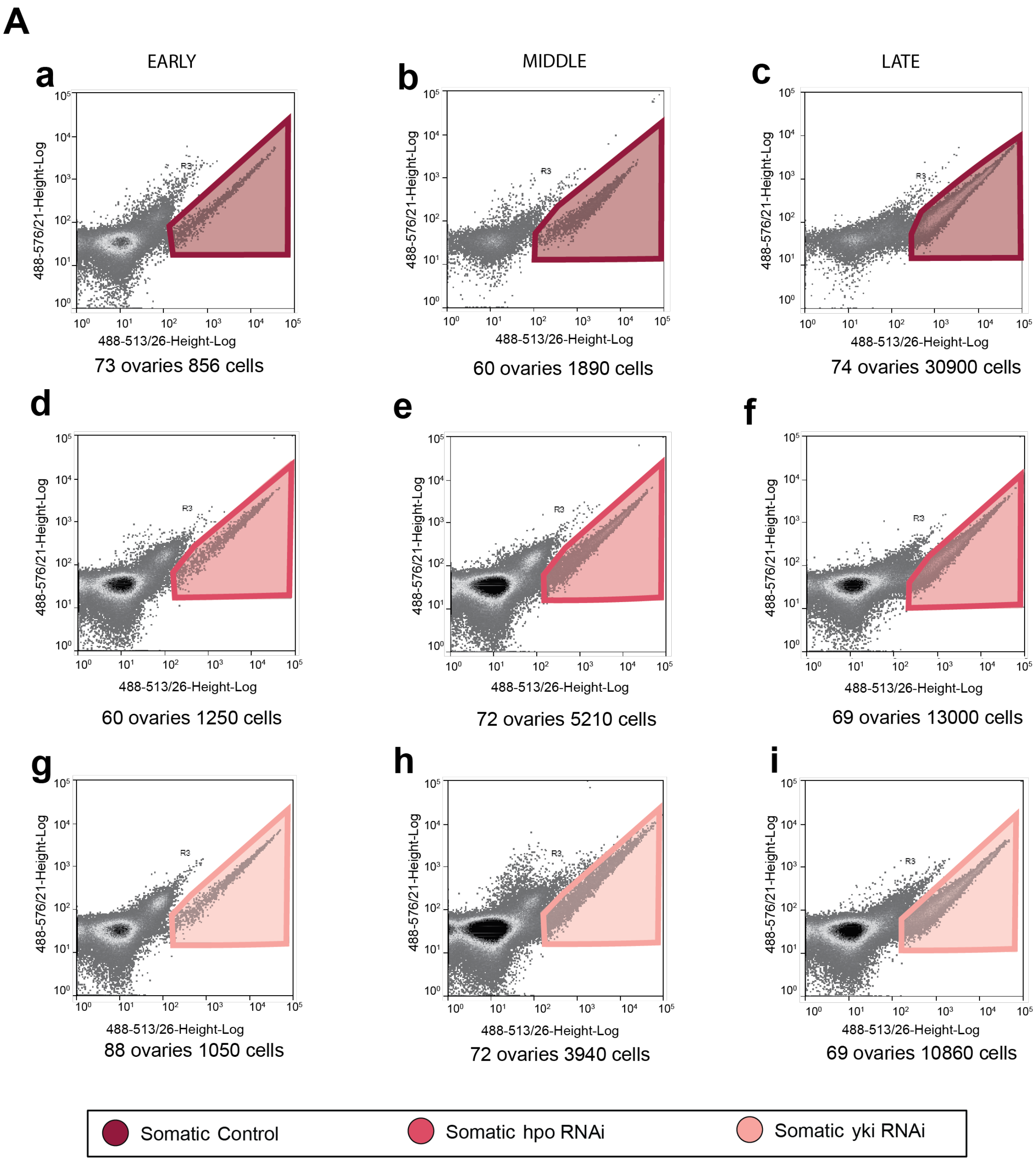
A
EARLY
MIDDLE
LATE
a
b
c
488-576/21-Height-Log
488-513/26-Height-Log
R3
73 ovaries 856 cells
60 ovaries 1890 cells
74 ovaries 30900 cells
d
e
f
60 ovaries 1250 cells
72 ovaries 5210 cells
69 ovaries 13000 cells
g
h
i
88 ovaries 1050 cells
72 ovaries 3940 cells
69 ovaries 10860 cells
Somatic Control
Somatic hpo RNAi
Somatic yki RNAi

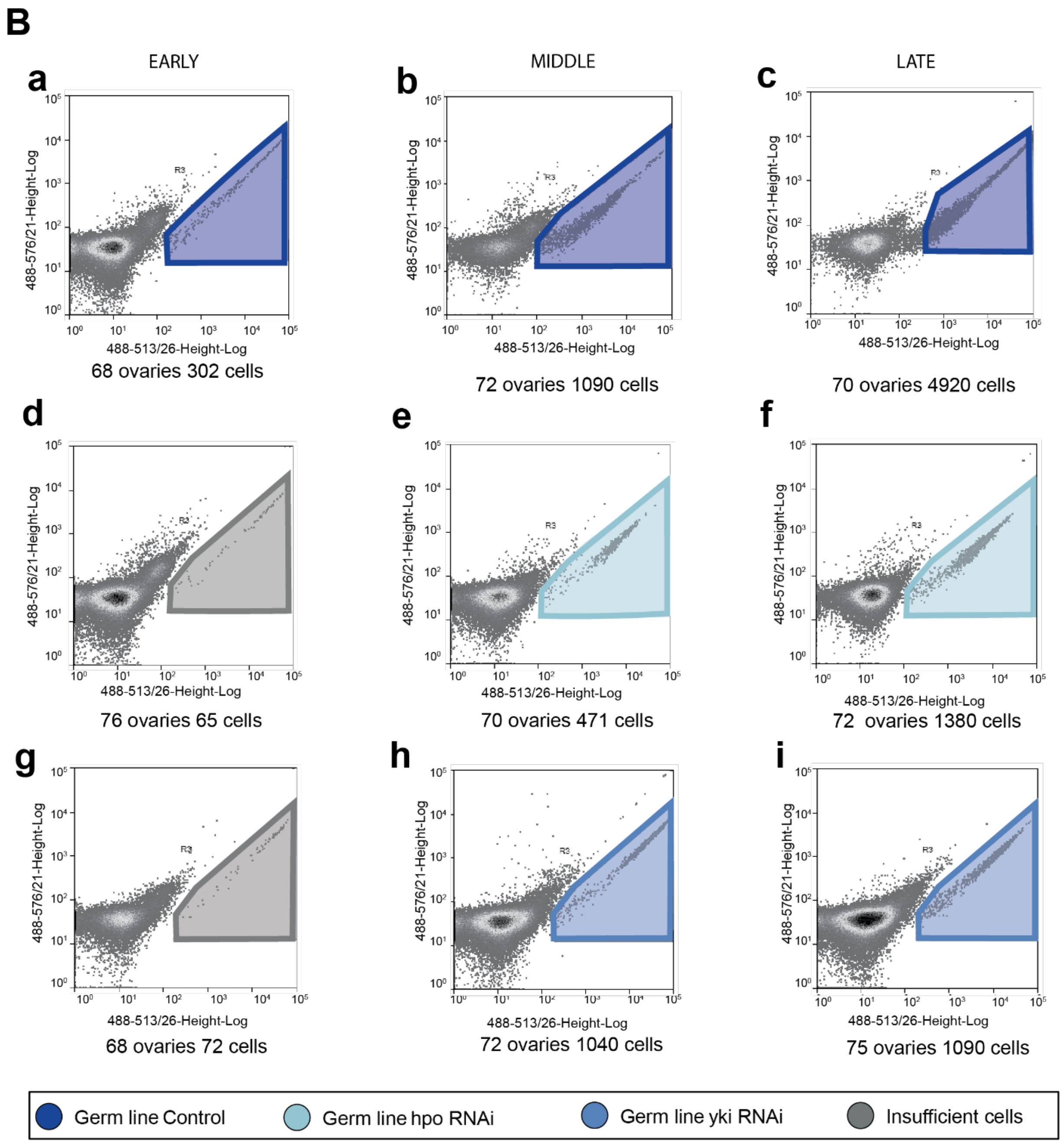


**Supplemental Figure S1: Representative FACS cell density plots. (A)** GFP-positive cell populations from dissociated control and RNAi treated- ovaries. GFP-positive somatic cells from early, middle and late stages are shown in three columns: 1 (panels a, d, g), 2 (panels b, e, h) and 3 (panels c, f, i). Number of ovaries used to generate cell pools and cell yields are shown below each plot. Representative plots show GFP-positive sorted somatic cells in (A-C) wild type control (D-F) *hpo* RNAi and (G-I) *yki* RNAi conditions. **(B)** GFP-positive germ line cells from early, middle and late stages are shown in three columns as in **A**. Early germ line cells under both *hpo* and *yki* knockdowns (gates colored in grey) yielded cell numbers sufficient for only a single replicate each and were not included in the final analysis.

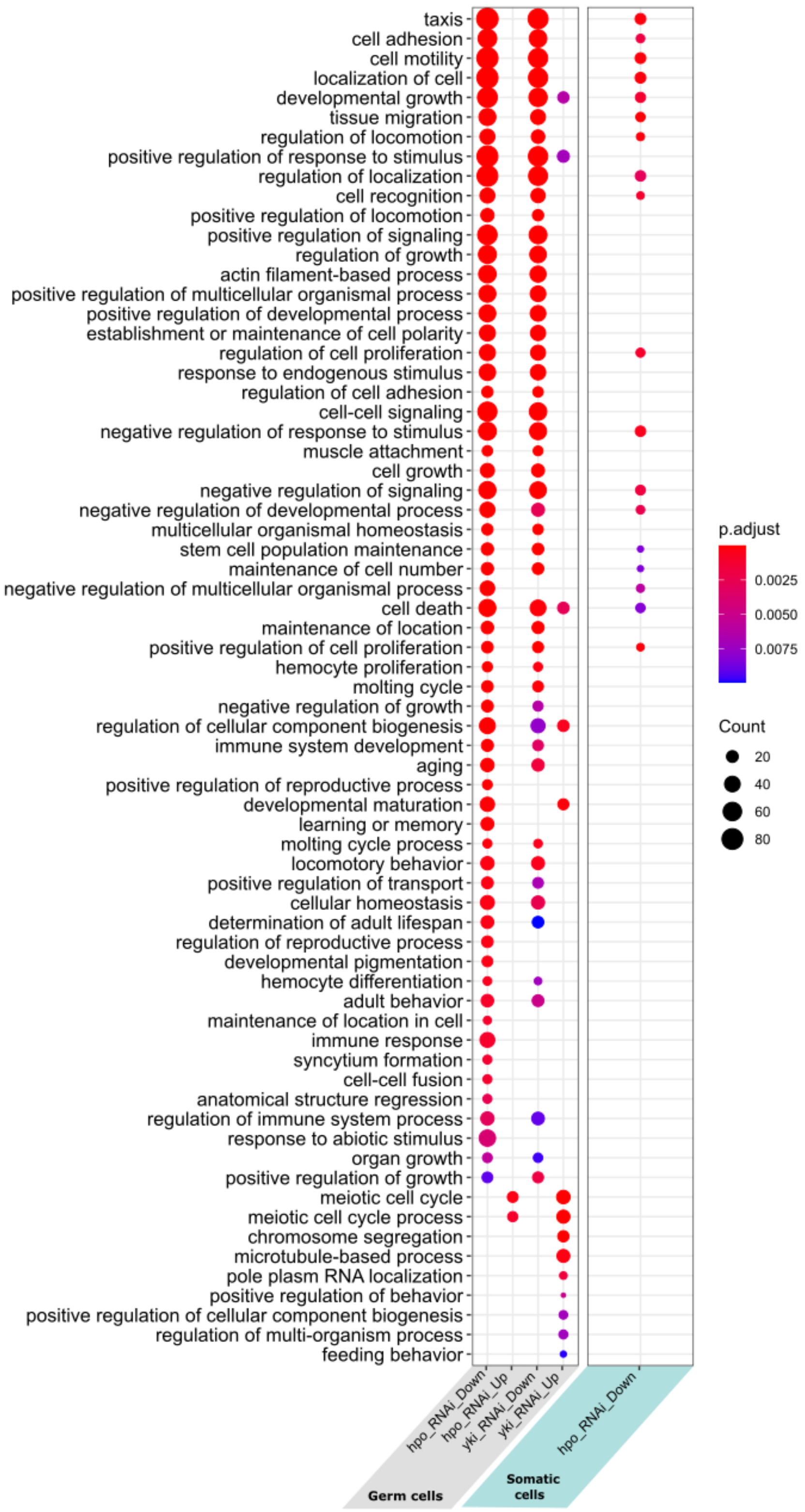


**Supplemental Figure S2: GO analysis.** Significantly enriched (Benjamini-Hochberg adjusted p-value<0.01 and q-value < 0.01) GO-terms of Biological Process level 3 within the significantly upregulated (Up) and downregulated (Down) genes in germ cells (left column) and somatic cells (right column) under *hpo* and *yki* RNAi conditions (grouping mid and late stages together).

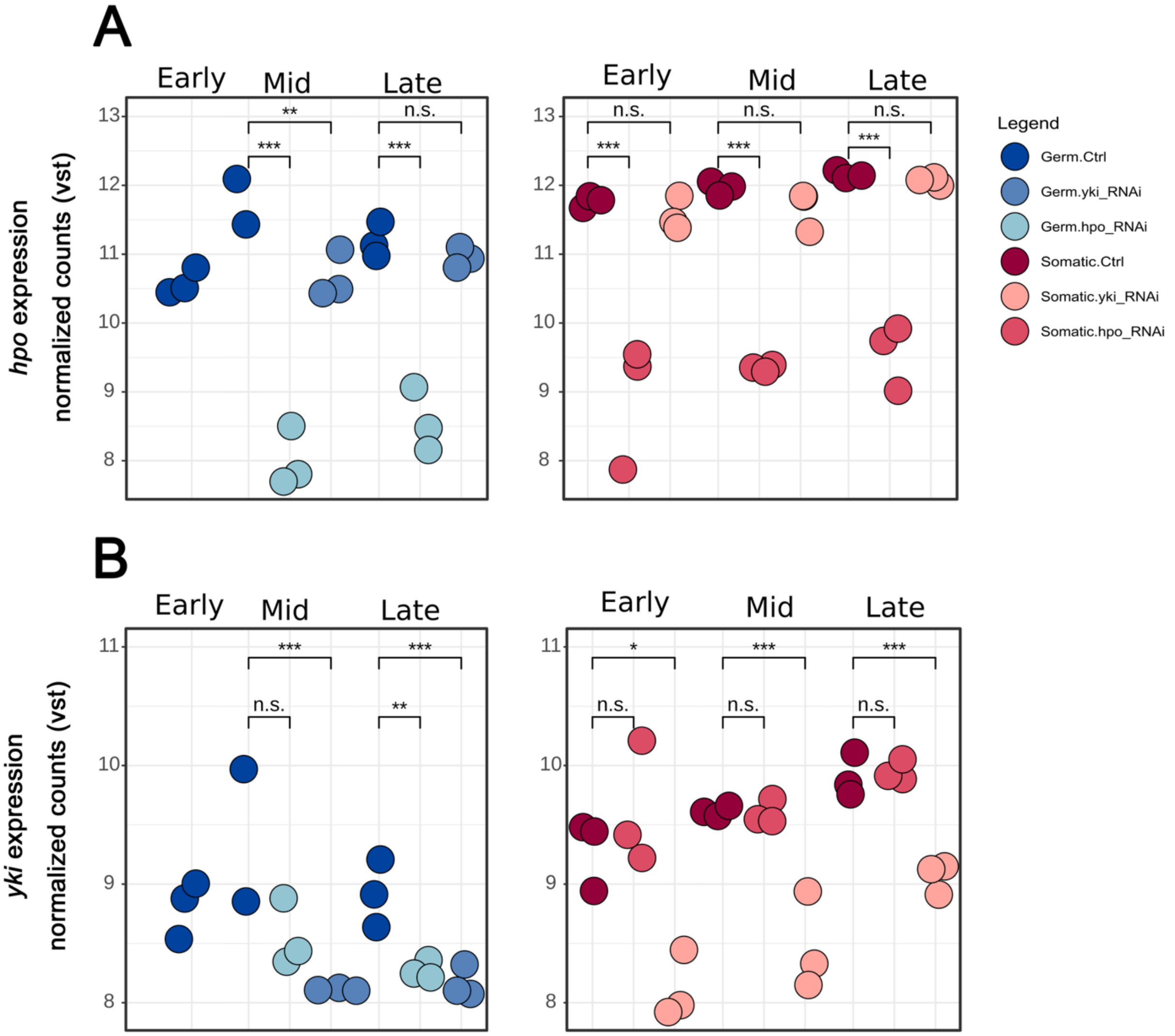


**Supplemental Figure S3: Expression levels of *hippo* and *yorkie* in the RNA-seq libraries after removing the mid stage germ cell control sample with anomalously low levels of Yki.** Expression levels of **(A)** *hippo* and **(B)** *yki* as vst-normalized counts in germ cells (left panels, blue dots) and somatic cells (right panels, red dots) in each RNA-seq library (color shades indicate genetic condition). The *hpo* and *yki* transcripts appear significantly downregulated in their corresponding RNAi conditions in each cell type. Additionally, *yki* appears significantly downregulated under *hpo* knockdown. Statistical significance levels from the differential expression analysis are codified as *** = adjusted p-value<0.01, ** = adjusted p-value<0.05, n.s. = not significant.

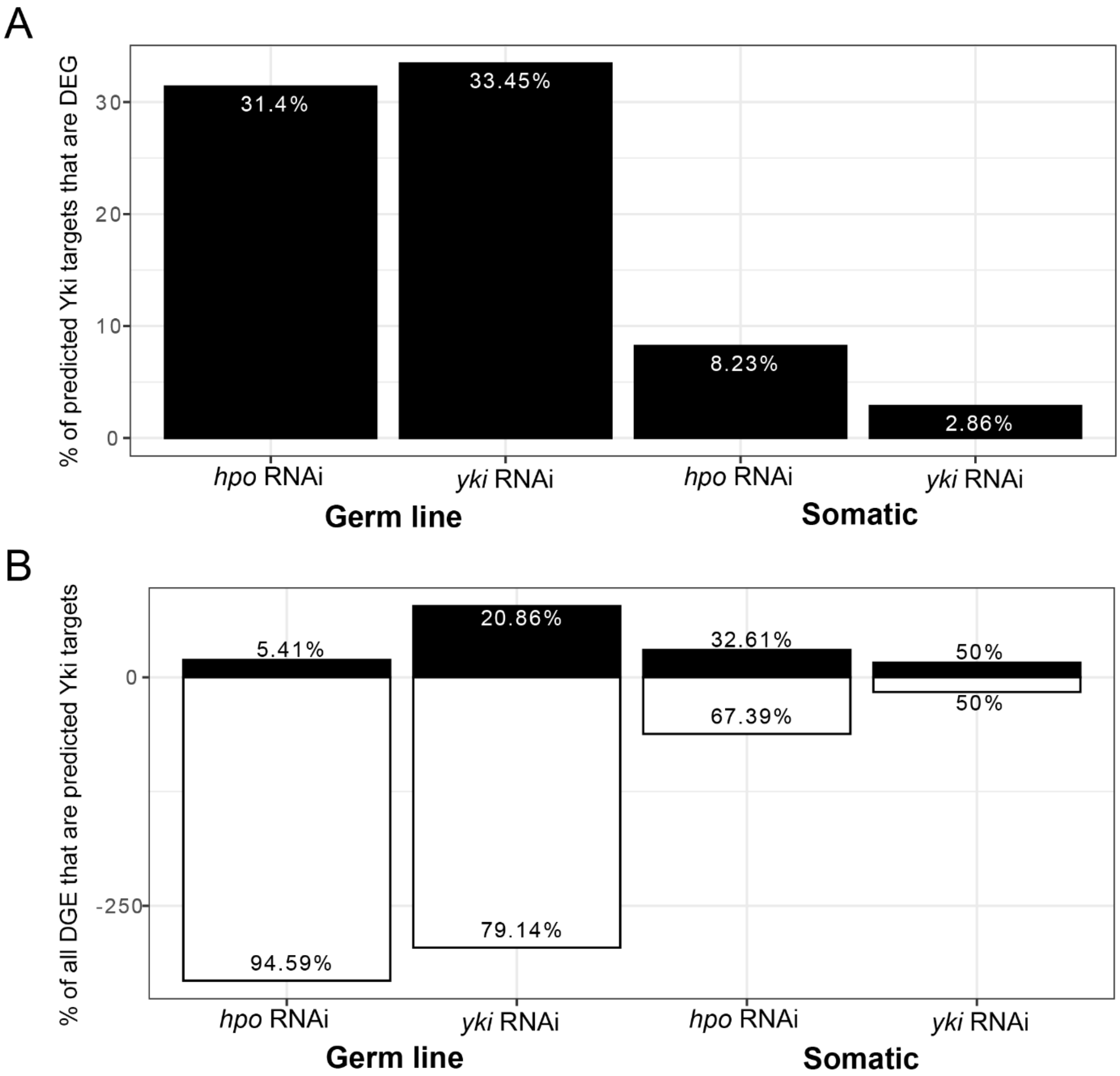


**Supplemental Figure S4: Change of expression levels of predicted direct Yki targets under knockdown conditions. (A)** Percentage of predicted direct Yki targets genes identified from published ChIP-seq data (Oh et al., 2013) that are significantly differentially expressed (adjusted p-value <0.05) under each knockdown condition (*hpo*[*RNAi]* and *yki*[*RNAi]*) in each cell type (somatic (left two categories) and germ line (right two categories)). **(B)** Percentage of predicted direct Yki targets genes identified from published ChIP-seq data (Oh et al., 2013) that are within the significantly differentially up- and down-regulated categories (adjusted p-value <0.05) in each knockdown condition (*hpo*[*RNAi]* and yki[RNAi]) in somatic cells (left two categories) and germ line cells (right two categories)).

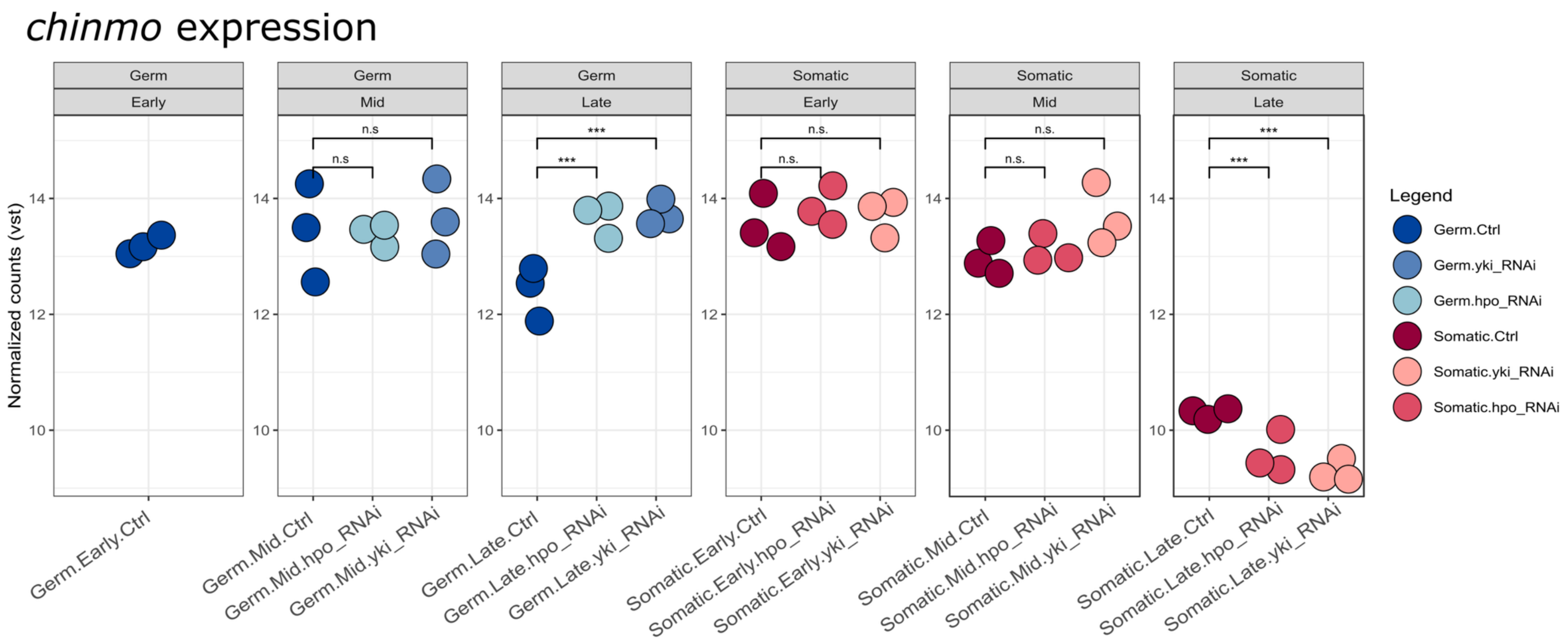


**Supplemental Figure S5: Expression of *chinmo* in each RNA-seq library.** The vst-normalized expression level of *chinmo* detected in each RNA-seq library (dots) grouped by cell type and stage, and colored based on cell type (blue: germ line, red: somatic) and genotype (color hue). The result of the differential expression analysis between conditions (control, *yki[RNAi]*, and *hpo[RNAi]*) within each cell type and stage is shown as the adjusted p-value with significance level indicators (n.s. = not significant and *** = padj <0.01) and log2 fold change.

A

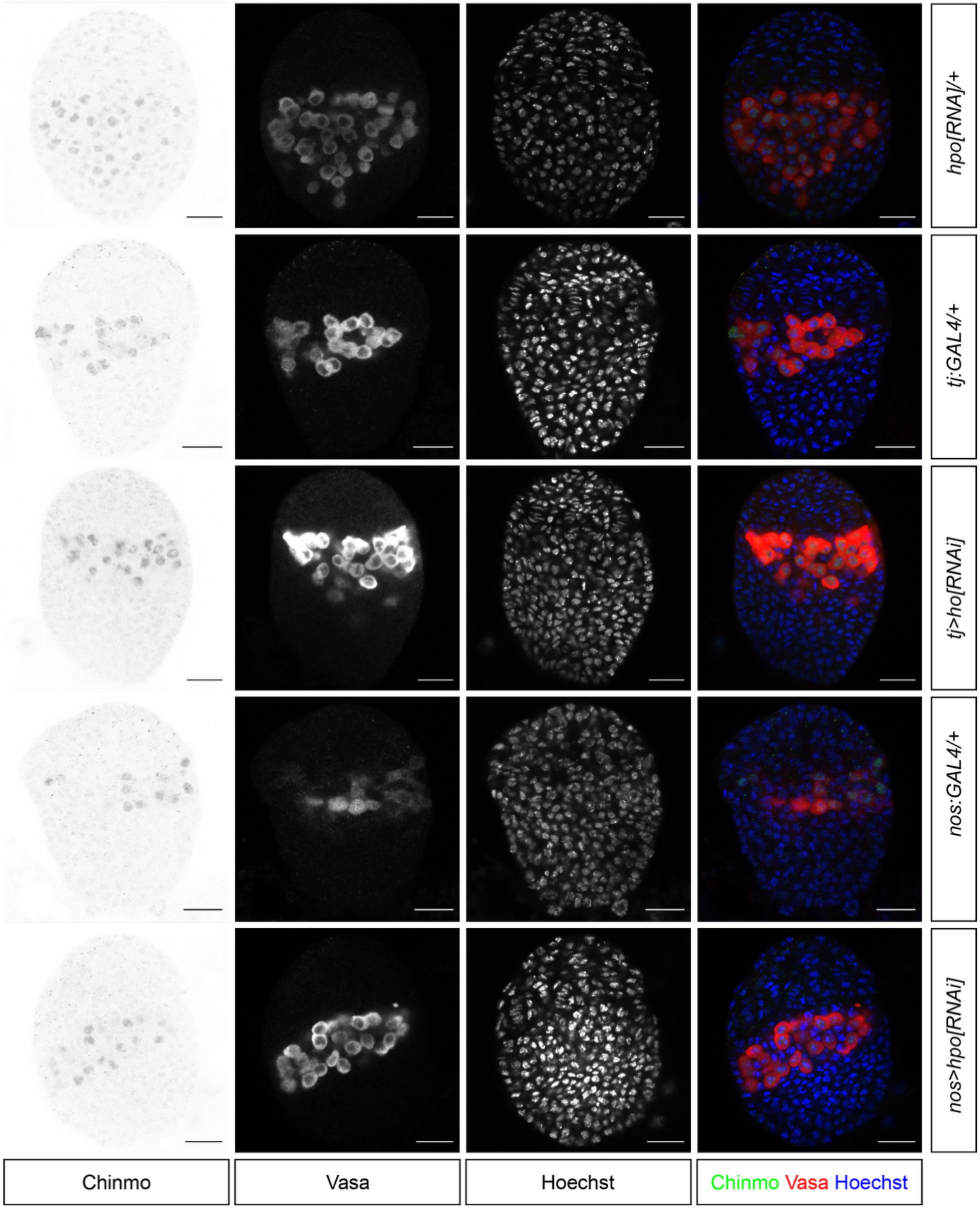
hpo[RNA]/+
tj:GAL4/+
tj>ho[RNAi]
nos:GAL4/+
nos>hpo[RNAi]
Chinmo
Vasa
Hoechst
Chinmo Vasa Hoechst

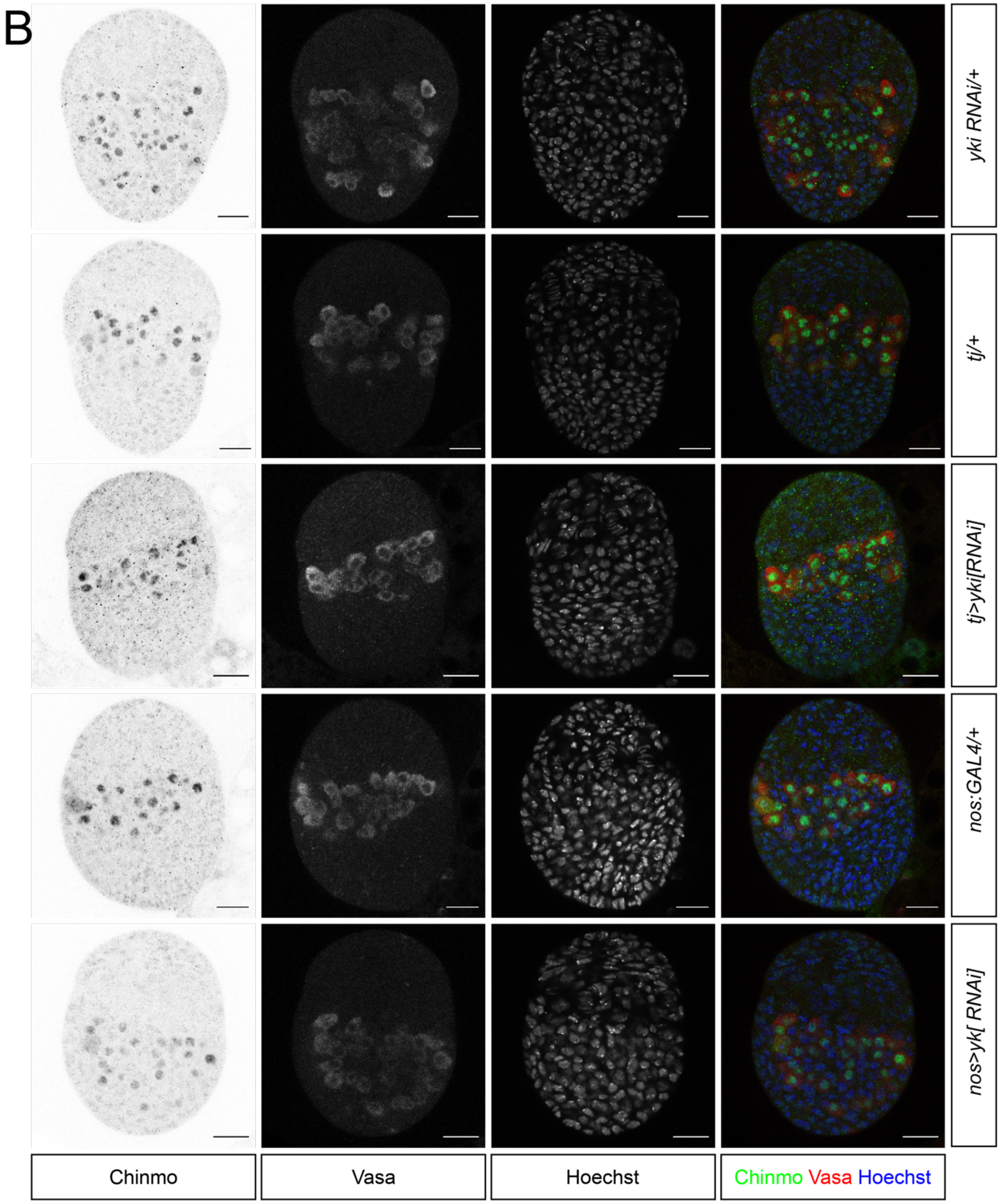
B
yki RNAi/+
tj/+
tj>yki[RNAi]
nos:GAL4/+
nos>yk[ RNAi]
Chinmo
Vasa
Hoechst
Chinmo Vasa Hoechst

C

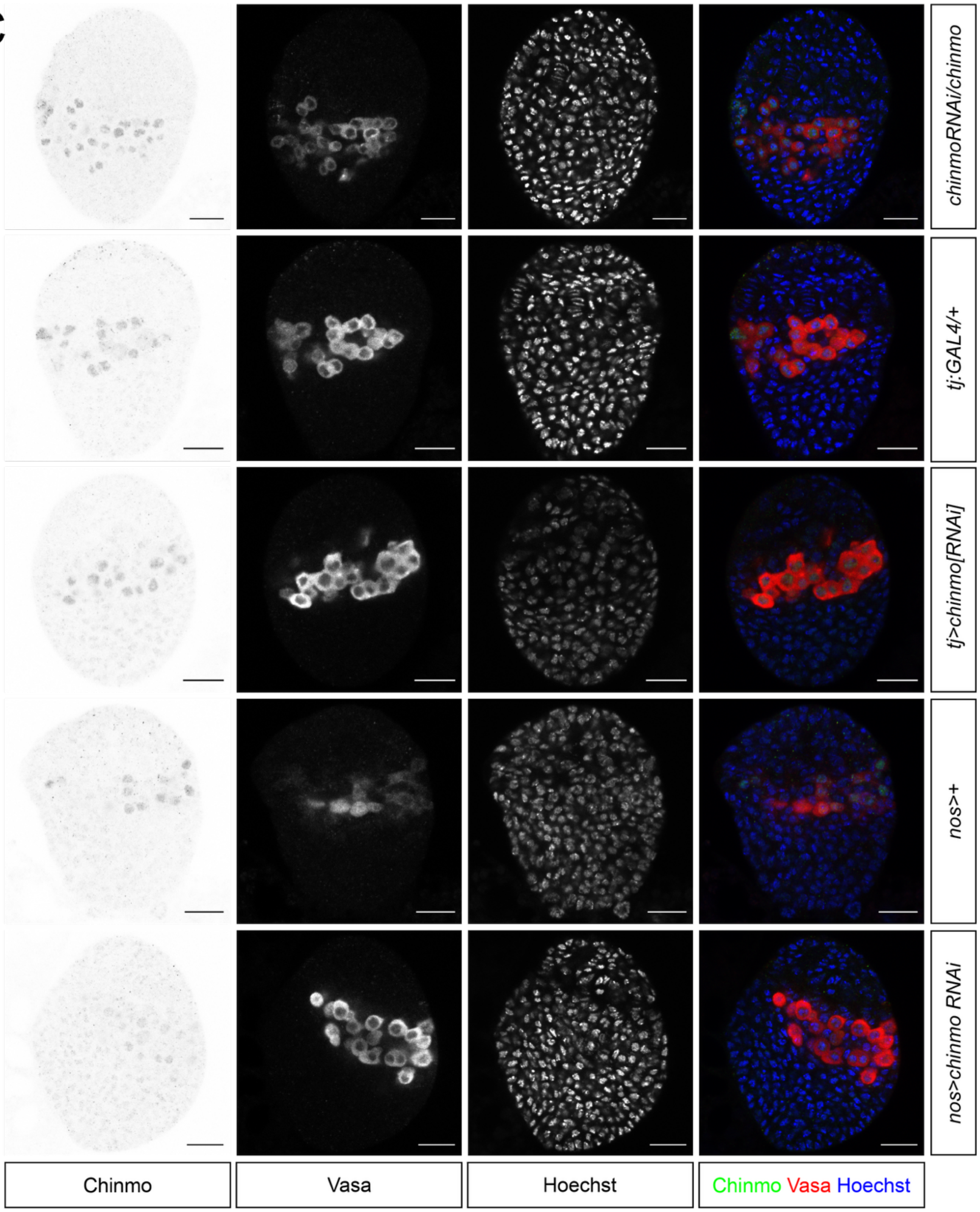
chinmoRNAi/chinmo
tj:GAL4/+
tj>chinmo[RNAi]
nos>+
nos>chinmo RNAi
Chinmo
Vasa
Hoechst
Chinmo Vasa Hoechst

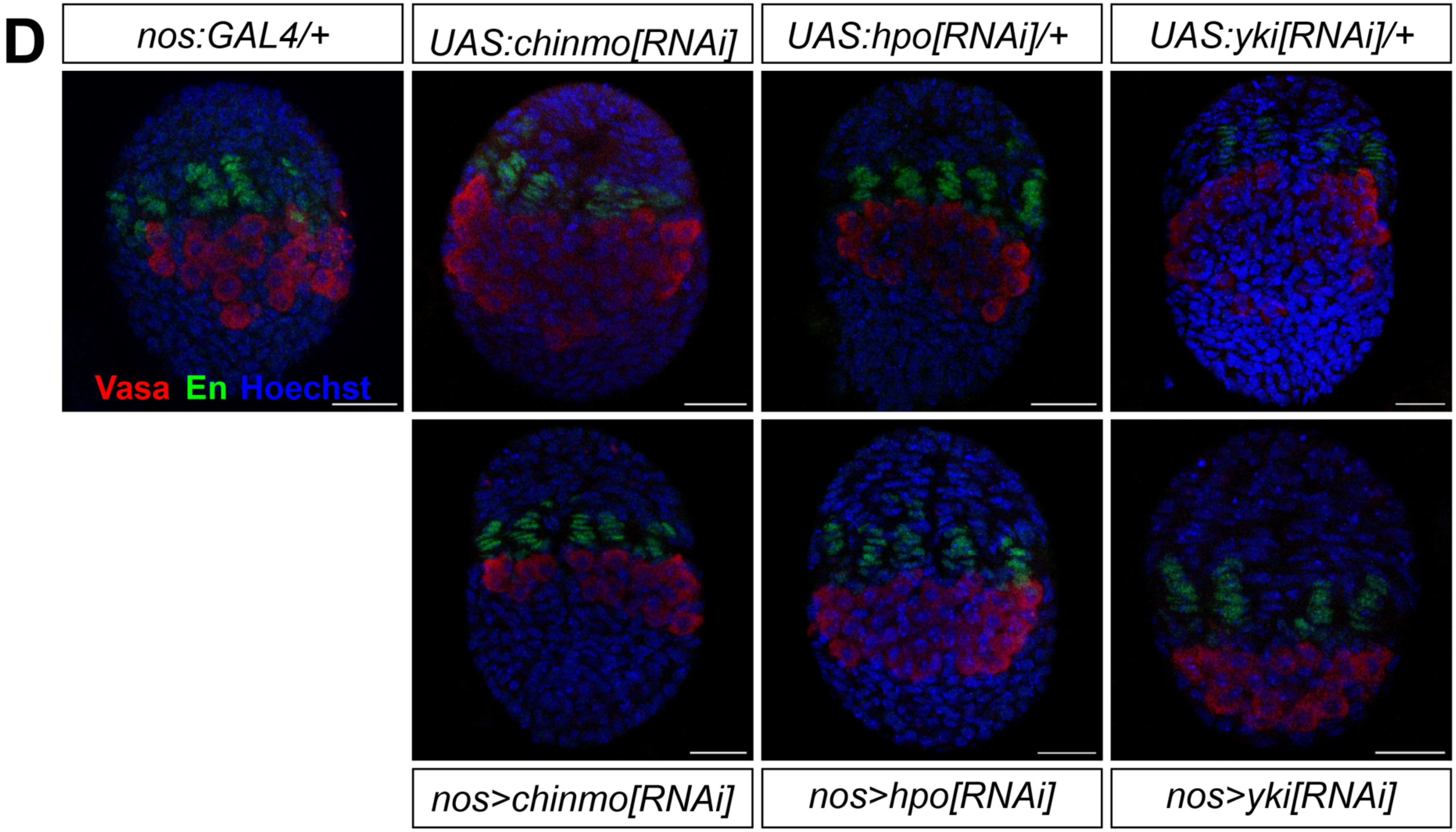


**Supplemental Figure S6: Detectable expression of Chinmo and Vasa proteins in larval ovaries of the genetic backgrounds analyzed in this study. (A)** Detection of Chinmo protein in the nuclei of germ cells under *hpo* RNAi driven in the somatic cells (by *tj:GAL4*) or in the germ cells (by *nos:GAL4*), and in appropriate controls. **(B)** Detection of Chinmo protein in the nuclei of germ cells under *yki*[*RNAi]* driven in the somatic cells (by *tj:GAL4*) or in the germ cells (by *nos:GAL4*), and in appropriate controls. **(C)** Detection of Chinmo protein in the nuclei of germ cells under *chinmo*[*RNAi]* driven in the somatic cells (by *tj:GAL4*) or in the germ cells (by *nos:GAL4*), and in appropriate controls. Red = Vasa; Chinmo = green, scale bars = 20μm. **(D)** Detection of Vasa protein in the nuclei of germ cells under *hpo*[*RNAi]*, *yki*[*RNAi]*, and *chinmo*[*RNAi]* driven in the germ cells (by *nos:GAL4*), and appropriate controls. Vasa staining was used to count germ cells in all conditions. Red = Vasa; Engrailed = green, scale bars = 20μm.

### SUPPLEMENTAL TABLE LEGENDS

**Supplemental Table S1:** File name ***Supplemental Table S1.xlsx***

**Replicates of tissue types and treatments.** Each row from left to right: Samples with description of the control or the treatment of tissue, biological replicates, technical replicates, stage of the ovary development, cDNA concentrations obtained from sorted cells, PrepX Index used in the sequencing cDNA library, number of ovaries pooled to prepare the replicate, total pooled FACS cell-counts from GFP positive cell-types, number of total raw sequenced reads, number of reads mapped to the genome, and the percentage of mapped reads.

**Supplemental Table S2:** File name ***Supplemental Table S2.csv***

**Principal Component Analysis and Gene list.** Values of principal component 1, 2 and 3 for all the differentially expressed genes followed by FlyBase gene ID, Gene symbol, gene name and FlyBase CG number.

**Supplemental Table S3:** File name ***Supplemental Table S3.csv***

**Differential gene expression.** List of genes and their state of regulation in somatic and germ line tissues. FlyBase gene ID, Log2 fold change in gene expression, Standard Error (SE) of Log2 fold change, stat (Wald statistic), p-value and adjusted p-value, upregulation and downregulation in *hpo*/*yki* RNAi conditions, ENTREZ gene ID, gene symbol, gene name, FlyBase CG gene ID and expression cell-type (somatic or germ cell).

**Supplemental Table S4:** File name ***Supplemental Table S4.xlsx***

**Tabulated data counts of quantified parameters in *chinmo* functional analyses.** Raw data collected for ovariole number, terminal filament-cell numbers, terminal filament numbers and fluorescence intensity levels of protein expression in the various genetic backgrounds examined.

**Supplemental Table S5**: File name ***Supplemental Table S5.xlsx***

**List of genetic crosses performed in this study**. List of fly lines used in this paper, crosses performed to generate knockdown conditions, and control crosses.
